# Var|Decrypt: a novel and user-friendly tool to explore and prioritize variants in whole-exome sequencing data

**DOI:** 10.1101/2022.09.02.506346

**Authors:** Mohammad Salma, Elina Alaterre, Jérôme Moreaux, Eric Soler

**Author notes:** correspondance : MS, ES.

## Abstract

**Motivation:** High throughput sequencing (HTS) offers unprecedented opportunities for the discovery of causative gene variants in multiple human disorders including cancers, and has revolutionized clinical diagnostics. However, despite more than a decade of use of HTS-based assays, extracting relevant functional information from whole exome sequencing (WES) data remains challenging, especially for non-specialists lacking in-depth bioinformatic skills.

**Results:** To address this limitation, we developed Var|Decrypt, a web-based tool designed to greatly facilitate WES data browsing and analysis. Var|Decrypt offers a wide range of gene and variant filtering possibilities, clustering and enrichment tools, providing an efficient way to derive patient-specific functional information and to prioritize gene variants for functional analyses. We applied Var|Decrypt on WES datasets of 10 acute erythroid leukemia patients, a rare and aggressive form of leukemia, and recovered known disease oncogenes in addition to novel putative drivers. We additionally benchmarked Var|Decrypt on an independent dataset of ~90 multiple myeloma WES, recapitulating the identified deregulated genes and pathways, showing the general applicability and versatility of Var|Decrypt for WES analysis.

## Introduction

Leukemia comprises a heterogeneous group of deadly blood cancers resulting from abnormal or impaired hematopoietic cell differentiation and stem cell function. Many cell intrinsic factors can contribute to leukemia initiation, development and maintenance, including mutations affecting signaling pathways, metabolic genes, splicing components and epigenetic regulators, leading to acquisition of several cancer hallmarks (Manolio *et al*., 2013). Although a number of recurrently mutated genes have already been identified in leukemia of both myeloid and lymphoid origins, a number of rare and/or aggressive leukemia subtypes, for which the driving oncogenes are poorly characterized, still require in depth analyses. This also applies to solid tumors and rare cancers types, for which the mutational landscape remains to be thoroughly characterized. In this context, high throughput sequencing (NGS) of patient samples is instrumental to unravel the underlying genetic abnormalities. The era of medicine of precision and customization – i.e. the capacity to provide patient care guided by genetic diagnostic, is at reach since genome-scale sequencing approaches such as whole exome sequencing (WES) are implemented in routine diagnostics (Manolio *et al*., 2013)(Xiao *et al*., 2021). WES offers a flexible and efficient way to highlight the mutational landscape of hundreds of patients at relatively low cost. Indeed, WES is mostly focused on the gene coding regions of the genome, representing ~2% of the total human genome sequence (approximately ~30 million base-pairs) (Bertier *et al*., 2016). Although WES is unable to highlight complex genomic rearrangements such as chromosomal translocations or inversions, it still provides highly relevant information regarding gene mutations, including splice site mutations. It is therefore extensively used in multiple clinical centers, and studies over the world (**Fig. 1**). As a growing number of patients became sequenced, increasing amounts of detected variants were published and added into public databases. The number of known disease genes has therefore dramatically increased over the past decade, which reinforces diagnostic test performance (Smith *et al*., 2019). Clinics in the USA (Jacob *et al*., 2013), France (Thevenon *et al*., 2016) and the Netherlands (Vrijenhoek *et al*., 2015) for instance report WES as a promising tool from the systematic use of NGS in patients. However, despite its extensive use and significant advantages, extracting relevant information from WES data still represents a challenge (e.g. identifying recurrently mutated genes from cohorts of patients, or the biological pathways which are significantly affected). WES data analysis requires advanced bioinformatics tools and skills, which prevents non-specialists such as wet lab scientists or clinicians from being able to navigate within the datasets and perform custom analyses. A great challenge for scientists and clinicians with limited or no bioinformatics skills is therefore to be able to manipulate WES data in order to extract biological meaningful information, and relevant genes or pathways to guide functional testing and therapeutic approaches.

**Figure 1.**
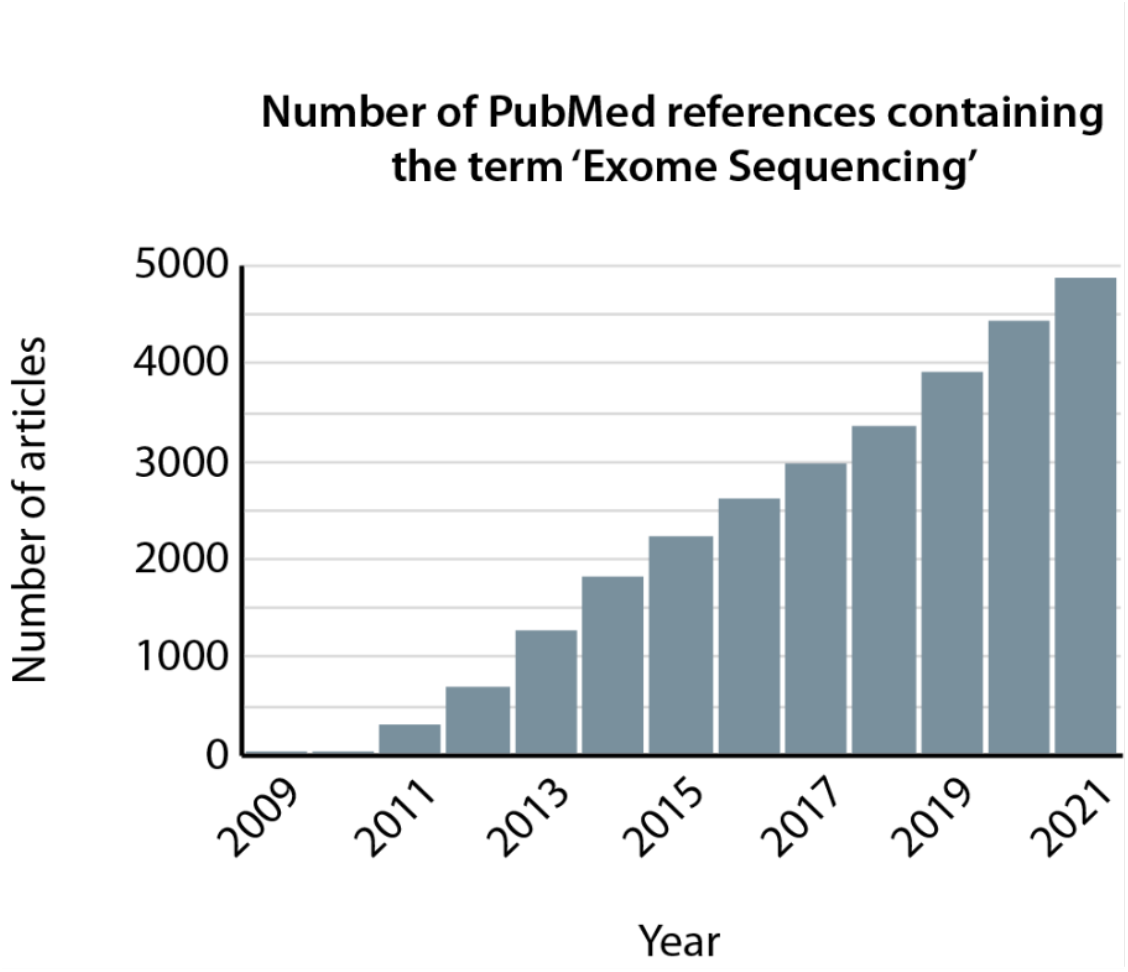
Widespread use of whole exome sequencing data. Graph plotting the number of PubMed articles containing the term 'exome sequencing', showing a continuous increase over the 2009-2021 period.

WES data analysis consists in two main features: first, the detection of variants or mutations, and second, the post variant calling analysis for prioritization of genes or variants within patient cohorts. The detection of germinal or somatic variants from matched tumor-control samples is still a complex task (Binatti *et al*.), despite the availability of dedicated tools. Recently, Bertier et al reported a total of 23 different challenges related mainly to the production, analysis and sharing of WES data (Bertier *et al*., 2016). They mentioned that the interpretation of variants of unknown significance was one of the most reported challenges across all articles. Often, the interpretation of this data requires specially trained staff (Bertier *et al*., 2016). In order to understand the biological meaning of these variants and/or differences between tumoral and control samples, two categories of prioritization tools exist. The first category focuses on variant annotation and filtering to eliminate those with low relevance such as SNPsift (Frontiers | Using Drosophila melanogaster as a Model for Genotoxic Chemical Mutational Studies with a New Program, SnpSift | Genetics), GEMINI (Paila *et al*., 2013) and VCF tools (Danecek *et al*., 2011), Ingenuity^®^ Variant Analysis™ Software (Home – QIAGEN Digital Insights), Golden Helix SNP & Variation Suite (SNP & Variation Suite (SVS) – Golden Helix), BiERapp (Alemán *et al*., 2014), EVA (Coutant *et al*., 2012), Exomiser (Smedley *et al*., 2015), Variant Ranker (Alexander *et al*., 2017), BrowseVCF (Salatino and Ramraj, 2017), TGex (Dahary *et al*., 2019) and VCF-Miner (Hart *et al*., 2016). The second category of prioritization tools attempts to establish a relationship between the detected variants and known diseases, pathways, biological process, etc. using various public databases. These include GeMSTONE (Chen *et al*., 2017) and BiERapp (Alemán *et al*., 2014). Other web-based tools like Enrichr (Chen *et al*., 2013), GOrilla (Eden *et al*., 2009), wKGGSeq (Li *et al*., 2015), geneontology (Gene Ontology Resource) and g:Profiler (Raudvere *et al*., 2019) can perform several enrichment analyses using a list of genes, which requires users to extract and prepare the input data according to the tool recommendations. In that respect, a flexible and user-friendly all-in-one solution, with built-in functionalities designed to facilitate extraction of relevant meaningful biological data from WES is clearly lacking. Here we present Var|Decrypt, a user-friendly and easy-to-use Rshiny application designed to fulfil this gap.

## RESULTS

### Exome-Seq analysis pipeline

To provide an all-in-one solution, we first implemented an Exome-seq variant analysis pipeline. The different steps are managed by Snakemake platform (Köster and Rahmann, 2012) which is a tool to create reproducible and scalable data analysis workflow (**Fig. 2**). It can be scaled to server, cluster, grid and cloud environment with no need to change the workflow definition [1]. The WES analysis pipeline contains all needed tools within a Singularity and docker container (see **Fig. 2**). It consists in the following steps: first, the reads are aligned to the human genome using BWA-MEM (Li, 2013) which is one of the most widely used well performing aligner for WES data. Post-mapping optimization is performed using GATK (McKenna *et al*., 2010; DePristo *et al*., 2011; Auwera *et al*., 2013; Poplin *et al*., 2018) suite which includes many tools to manage WES and Whole genome sequencing (WGS) data. For the variant calling step, the HaplotypeCaller is used for germline variants and Mutect2 for somatic ones from GATK4 (Poplin *et al*., 2018) as recent studies have shown the efficiency of these tools in different scenarios (Cornish and Guda, 2015; Hwang *et al*., 2015)(Xiao *et al*., 2021). After the variant calling procedure, the different variants need to be subsequently annotated. To this aim, we used the popular and powerful ANNOVAR tool (Wang *et al*., 2010) which integrates several useful databases (for example, dbSPN, 1000 genomes and Clinvar annotations databases). Our pipeline comes with a python master script (run_preVarDecrypt.py) which allows users to run the pipelines’ steps using one command line. Its requires two config files; 1) first one containing the path to fastq files and some optional parameters, 2) the second is tabulated file to make the association between tumoral and control samples.

**Figure 2.**
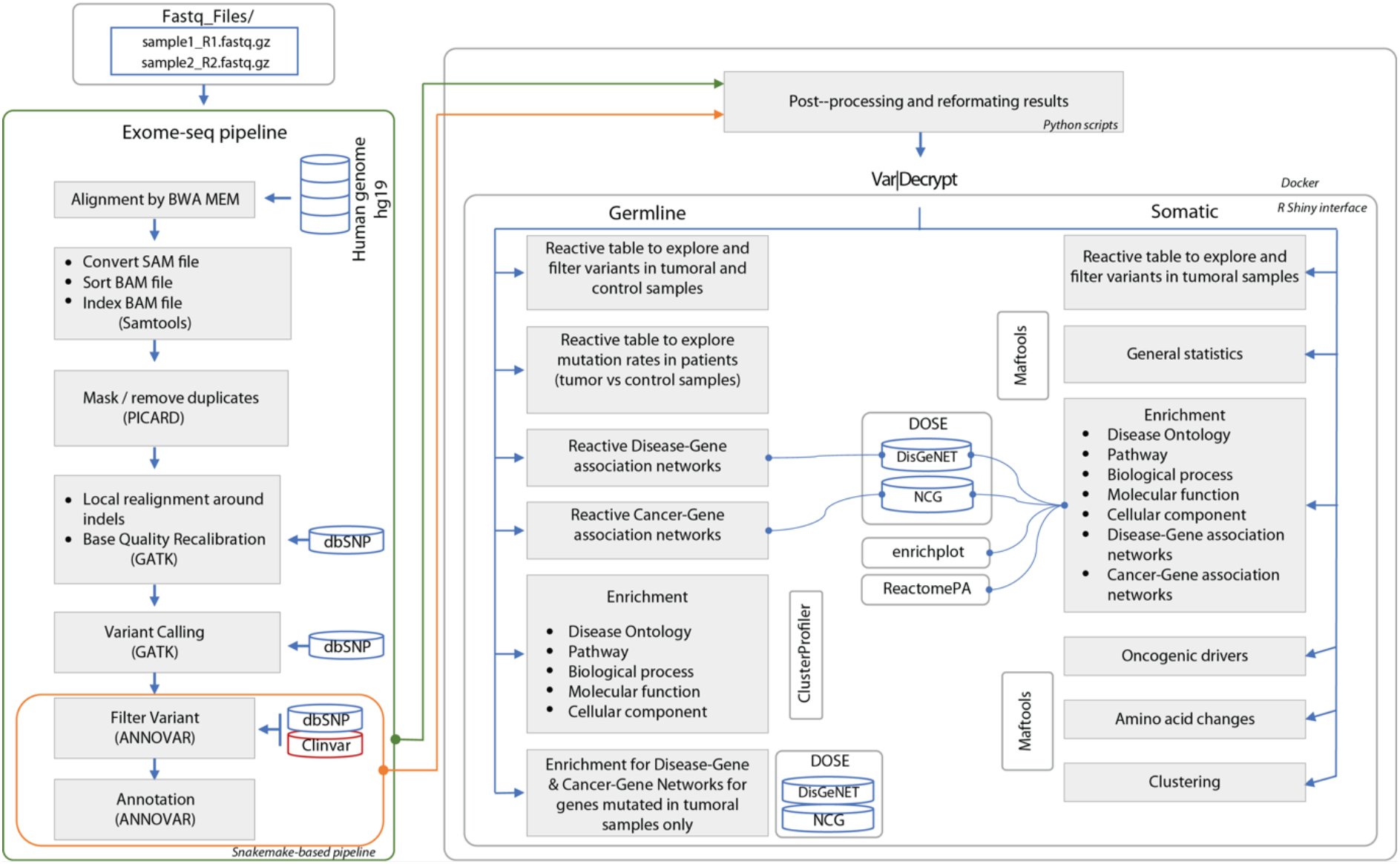
Bioinformatic pipelines used to process whole exome sequencing data. The WES and Var|Decrypt pipelines are depicted on the left and right, respectively. The various filtering steps and packages used are indicated. Two versions of the WES pipeline are available (The part highlighted in green corresponds to the pipeline to process WES data starting from fastq files; in orange the one allowing users to directly processing vcf files).

### WES data post-processing using Var|Decrypt

In order to facilitate WES data analysis and functional interpretation, we developed Var|Decrypt, an easy-to-use and user-friendly R Shiny interface. It includes several R packages to perform different post-VCF downstream analyses (**Table 1**), which usually require users with scripting skills to perform tasks such as installing packages, preparing the input data and calling the appropriate function. Var|Decrypt imports the output results from the Exome-seq pipeline and provides many built-in enrichment analyses options, helping researchers to develop or confirm hypotheses, to easily explore the differences between normal and tumor samples, and to prioritize variants, genes and pathways for functional analyses. Var|Decrypt is a fast-operating tool which provides multiple outputs within short time frames (i.e. seconds to minutes for loading a full dataset). The output results and variables are saved in an Rdata file which lets users to explore Var|Decrypt results subsequently, instead of re-running the analysis. Var|Decrypt allows to explore, filter, sort genes containing variants, or to search for a specific gene through dynamic interfaces (see below).

**Table 1.** Comparison of Var I Decrypt with commonly used enrichment analysis tools. The table depicts the various advantages of Var I Decrypt over the main online tools for gene ontology and biological feature enrichment analyses. (1) only kegg enrichment analysis needs an internet connection (2) control can be used only as background

| Name | Var\|Decrypt | GeMSTONE | GOrilla | geneontology | gprofiler/2 | wKGGSeq | Enrichr |
| --- | --- | --- | --- | --- | --- | --- | --- |
| **Variant type** |  |  |  |  |  |  |  |
| Germline | Yes | Yes | - | - | - | - | - |
| Somatic | Yes | No | - | - | - | - | - |
| **Enrichment analysis** |  |  |  |  |  |  |  |
| Input Files | post_processing files | Internal format | List of genes | List of genes | List of genes | VCF | List of genes |
| Save output results | Yes | Yes | No | Yes | Yes | Yes | Yes |
| Explore and filter variants by gene and by patient | Yes | No | No | No | No | No | No |
| Gene mutation rate in the cohort | Yes | - | No | No | No | No | No |
| Dynamic exploration | Yes | No | No | - | - | No | No |
| Independent output table by enrichment | Yes | No | Yes | Yes | Yes | Pathway only | Yes |
| All involved genes | Yes | No | Yes | Yes | Yes | - | Yes |
| Compare Tumoral/control | Yes | No | No<sup>(2)</sup> | No | No | No | No |
| **Visualization** |  |  |  |  |  |  |  |
| General statistics | Somatic only | Yes | No | No | No | Yes | No |
| Dynamic enrichment plots | Yes | No | No | No | No | No | Yes |
| Customizable | Yes | No | No | No | No | No | No |
| **Use** |  |  |  |  |  |  |  |
| User-friendly interface | Yes | Yes | Yes | Yes | Yes | Yes | Yes |
| Programming skills | No need | No need | No need | No need | No need/Yes | No need | No need |
| Run locally | Yes | No | No | No | No/Yes | No | No |
| Run offline | Yes<sup>(1)</sup> | No | No | No | No | No | No |

### Overall presentation of Var|Decrypt

The Var|Decrypt interface is composed of several tabs that each allows users to get a general overview of the Exome-seq data and allow to browse the mutated gene lists or focus on single genes, single variant types (e.g. stop gain, frameshift deletions, etc.).

The ‘Somatic variants explorer’ tab provides a gene list and summary table containing all detected mutated genes (we define the somatic variants as being the ones specifically acquired in the tumor sample as compared to the control cells; variants or mutations present in the control cells are considered as germline variants as they are not somatically acquired). (**Supp. Fig.1**). For each gene in the table, the total number of variants detected is indicated, together with the different types of variants identified, and the percentage of patients bearing a mutation in a particular gene. The right part of the table shows for each gene which patient sample contains the indicated variants (see **Supplementary Table 1** as an example). Instead of focusing on the variants themselves, this dynamic table is gene-centered, and it also provides information on the number of variants detected in the cohort for each gene, the types of variants (e.g. stop gain, frameshift deletions, etc.) and the percentage of patients bearing mutations on a particular gene. When using the ‘mutation rate’ column, users can sort the entire mutated gene list by mutation frequency (i.e. number of patients showing a mutation or variant within a given gene), which provides an overview of the top mutated genes. All types of variants are shown by default, but users may choose to highlight only a subcategory of variants such as stop gain, frameshift variants (deletions, insertions), etc. Whereas the germline variants from a patient are usually used to filter-out nonspecific variants in cancer samples, Var|Decrypt also allows working on the germline variants (‘Germline variant explorer’) which is useful for the study of mendelian genetic disorders or family case studies (not shown here).

The ‘General statistics’ tab provides information on the frequency of variant types within the cohort using a color code for the different types (e.g. frameshifts, non-sense, missense, etc.), the class of SNV (e.g. C>T, T>G, etc.) which may be useful to check if a particular bias is present in the samples or in the disease under study (**Fig. 3**). This tab also provides information on the total number of variants per sample, a feature that helps to quickly spot any outlier within the datasets. As exemplified in **Fig. 3**, sample m_13_D from our cohort contains ~30 fold more variants than the average of the other samples, likely arising from technical issues during the sequencing or sample handling procedure. Such problematic samples can therefore be quickly spotted and excluded from further analyses. Finally, the top 20 mutated genes are shown with the same color code as for the variant types, to get an overview of the recurrently mutated genes (**Fig. 3**).

**Figure 3.**
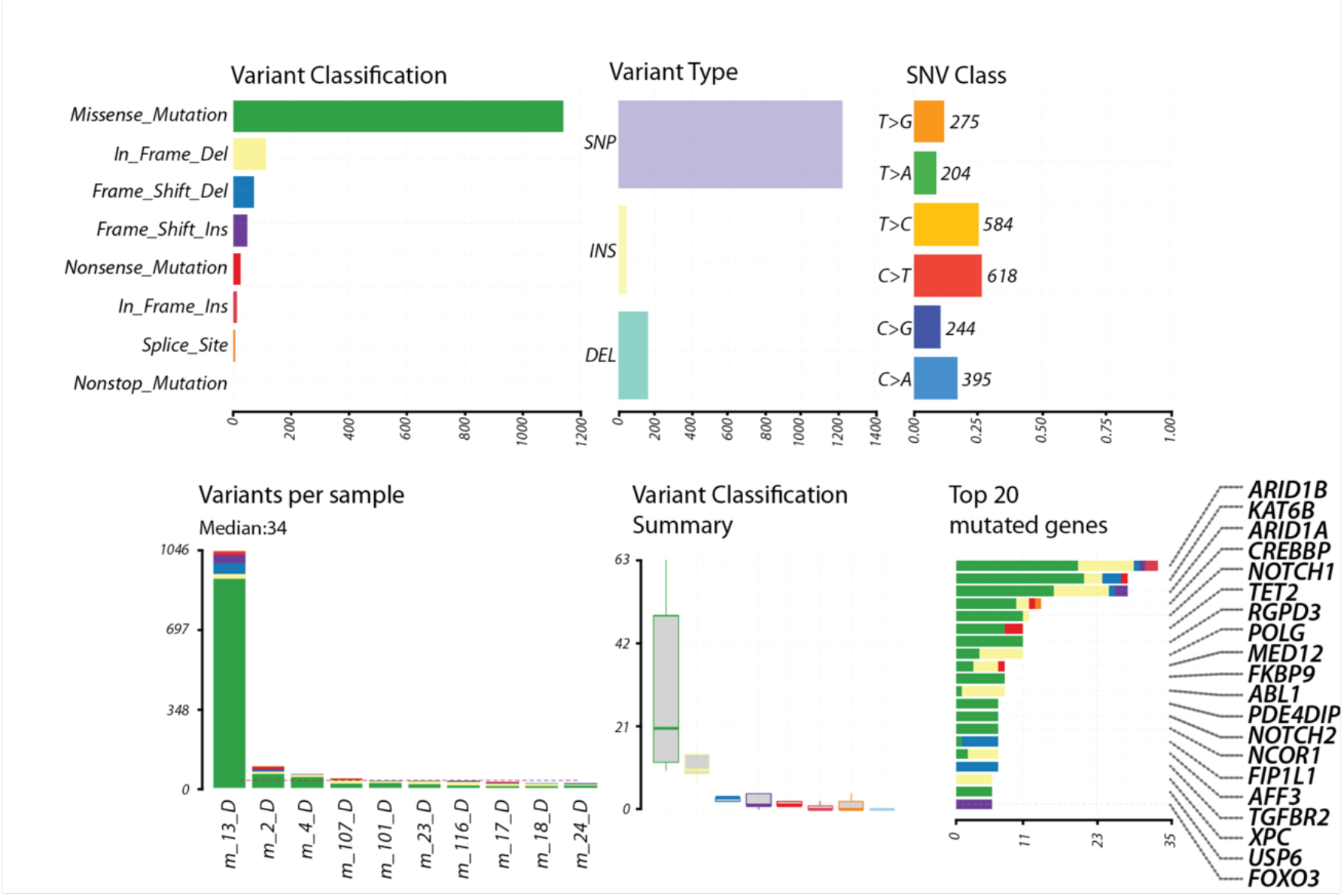
General overview of the WES datasets. The general features often erythroleukemic samples are displayed, showing the variant classification (color-coded as a function of the type of mutation, top left), variant type (single nucleotide polymorphisms (SNP), insertions (INS) and deletions (DEL), top middle), and single nucleotide variant (SNV) class (top right). The bottom panel displays the number of variants per sample (each column represents a unique patient), using the same color code as in the variant classification panel. Note that patient m_13_D is spotted as being an outlier with ~30 fold more variants than in the other patients. The dashed red line represents the median number of variants in the cohort. The middle panel shows the variant classification summary in the cohort, using the same mutation-specific color code. Finally, the bottom right panel shows the top 20 mutated genes in the patient cohort (the number of variants/mutations is shown on the horizontal axis), with the percentage of patients bearing a mutation in a given gene indicated.

### Identifying the recurrently mutated gene fraction within a patient cohort

Var|Decrypt offers the opportunity to quickly and easily browse WES data in order to identify recurrently mutated genes. By navigating in the somatic menus, users can in one click access the gene mutations frequencies (i.e. gene mutation percentage within the cohort), an important feature allowing to point at key genes likely involved in the disease phenotype. One key step in the discovery of cancer drivers is to be able to pinpoint the recurrently mutated genes within patient cohorts, as recurrently mutated genes likely represent true oncogenic drivers or genes important to sustain the cells’ transformed state. However, despite all the filtering steps applied in various Exome-Seq analysis pipelines, a very large number of variants usually remains, especially in cancer samples. This represents one of the main challenges to prioritize gene mutations when dealing with Exome-Seq datasets.

### Filtering of putative false positive gene mutations

A common issue of Exome-Seq data from short reads-associated sequencing platforms (such as Illumina sequencing) is the large fraction of variants called at genes harboring repetitive sequences, such as variable number of tandem repeats (VNTRs). The MUC gene family (Debailleul *et al*., 1998) is a good example of such problematic alignment and variant calling situation, as they contain long polymorphic stretches of ~60bp repeats VNTRs, which is problematic with the current aligners and variant callers. Although some true causative variants may indeed be present within the VNTRs of the MUC gene family (Kirby *et al*., 2013), we created a filtering option allowing users to define a threshold for the maximum number of variant allowed per gene, in order to ‘clean-up’ the mutated gene list and get rid of the error-prone VNTR-containing gene sequences in the patient cohort. As a result, by setting a threshold of a maximum of 4 variants per gene in a maximum of 20% of the patients, we could get rid of the apparently highly variable and likely false positive mutated genes in the final list (**Fig. 4**).

**Figure 4.**
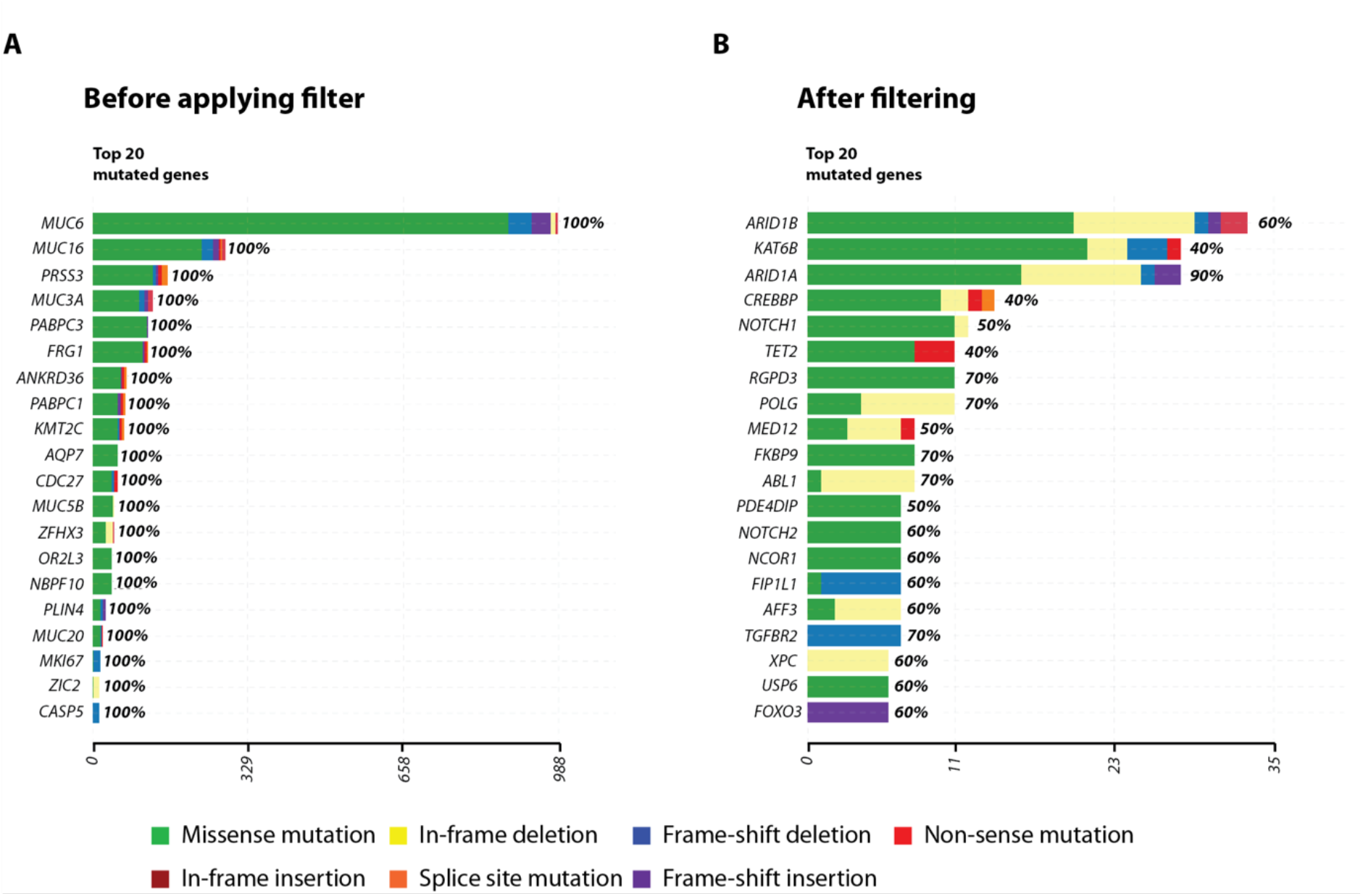
Custom filtering of putative false positive mutations. The top 20 mutated genes are show before (A) and after (B) applying the custom filters. This shows that without this filtering step, a number of genes score positive in 100% of the patients, including gene families containing variable number of tandem repeats (e.g. the MUC gene family). After applying a threshold (maximum of 4 variants per gene in a single patient, in a maximum of 20% of the patients) and selecting the option to retain genes present in the COSMIC database, the resulting mutated gene list is highly enriched in known oncogenic drivers and previously reported AEL mutated genes.

Another commonly used strategy to enrich for putative causative variants is to filter the mutated gene lists against cancer gene databases such as COSMIC, OncoKB or NCG (Vikova *et al*., 2019; Iacobucci *et al*., 2019a; Fagnan *et al*., 2020). We also provide a filtering option allowing to focus on the mutated genes that are tagged as cancer-associated from such databases. The resulting outputs therefore are highly enriched in putative oncogenic drivers, allowing to explore the mutational landscape of human cancers. As confirmation, applying such filtering strategy on our AEL WES data produced a mutated gene list enriched for previously reported AEL-associated gene mutations (Iacobucci *et al*., 2019a; Fagnan *et al*., 2020) such as the epigenetic modifiers TET2, NCOR1, NCOR2, BCOR, BCORL1, the CBP(*CREBBP*)/p300(*EP300*) co-activators, the polycomb repressive complex proteins EZH2, ASXL1, ASXL2, and the cohesin complex component RAD21 (**Supp Table 1**).

### Integration of enrichment tools

An important aspect of Var|Decrypt is the access to various types of enrichment analyses thanks to the implementation of dynamic customizable graphical outputs. Var|Decrypt contains different disease ontology, gene ontology (e.g. biological process, molecular function, cellular component), and Reactome/Kegg pathway enrichment tab offering the opportunity to identify particular pathway of functional alterations in the samples. The ‘enrichment’ tab offers users to quickly identify enrichments of disease ontology terms, biological pathways (Reactome, KEGG and WIKIpathways), or Gene-Ontology (GO)-terms such as ‘Biological Process’, Molecular Function’, or ‘Cellular Component’ linked to the mutated gene lists. In addition, searches for Gene-Disease associations or Gene-Cancer associations are also available to highlight many known associations with established human disorders and cancers. For each category, users can choose between three different graphical outputs including bar graphs, association or enrichment factor along with color-coded p-value representations (**Fig.5 A-C**). These outputs are dynamic and customizable as users can switch from one representation to another or increase/decrease the number of categories to display in one click. Var|Decrypt also provides a somatic interaction view in order to identify which gene mutations tend to co-occur or are mutually exclusive. In the example shown in **Fig. 5D**, BCOR and XPC mutations seem to be mutually exclusive, suggesting that inhibiting BCOR activity in XPC mutated AEL cells (and vice versa) may be therapeutically beneficial.

**Figure 5.**
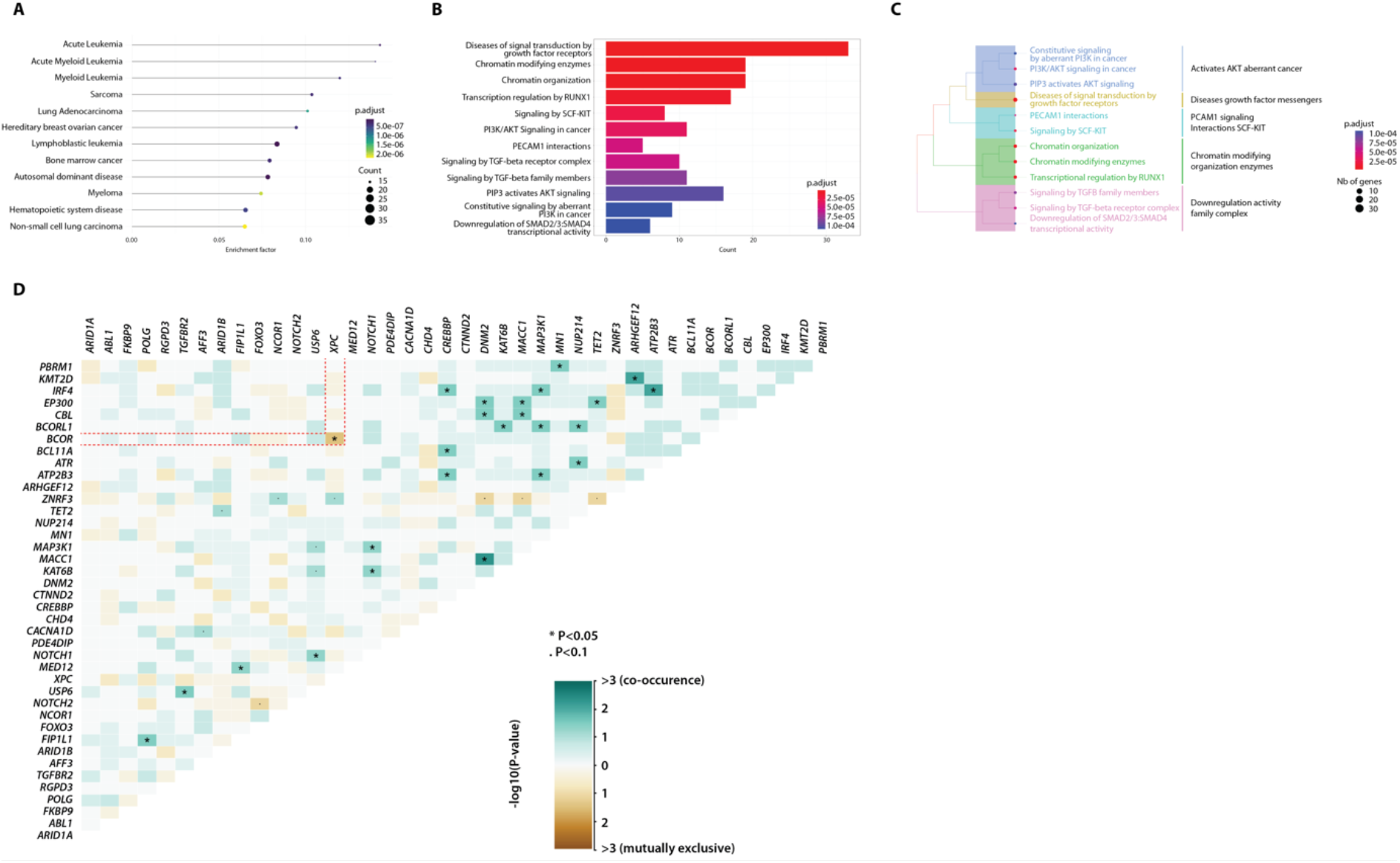
Disease and pathway enrichment features. Var|Decrypt allows to depict various enrichment plots using enrichment factor (A), qValue bar plot (B) or cluster tree (C) visualization for disease ontology, biological pathways (Reactome, Wiki, KEGG), various gene ontology (GO) categories (biological process, molecular function, cellular component), gene-disease and gene-cancer associations. (D) Matrix showing the mutually exclusive (brown) or cooperating mutations (green) in the AEL patient cohort. Dashed red lines highlight the mutually exclusive BCOR and XPC mutations.

Another useful built-in feature is the enrichment of mutations in genes belonging to known oncogenic signaling pathways. This feature provides a graph representation of the enriched mutated pathways together with the number of patients bearing mutations in the related pathways (**Fig.6A**). By simply clicking on a given pathway (right part), users can display a detailed list of the genes contained in the chosen pathway and check which gene and which patient sample harbor the mutation(s). This representation is useful to identify the recurrently mutated genes within a single oncogenic pathway and to check which signaling component is frequently altered in the disease. The example depicted in **Fig.6B** shows that the Receptor tyrosine kinase/RAS pathway, and the Notch and TGF-β pathways are frequently altered in AEL patients, and that the ABL Proto-Oncogene 1 (*ABL1*), *NOTCH2*, and TGF-β receptor 2 (*TGFBR2*) receptor genes are among the top mutated genes.

**Figure 6.**
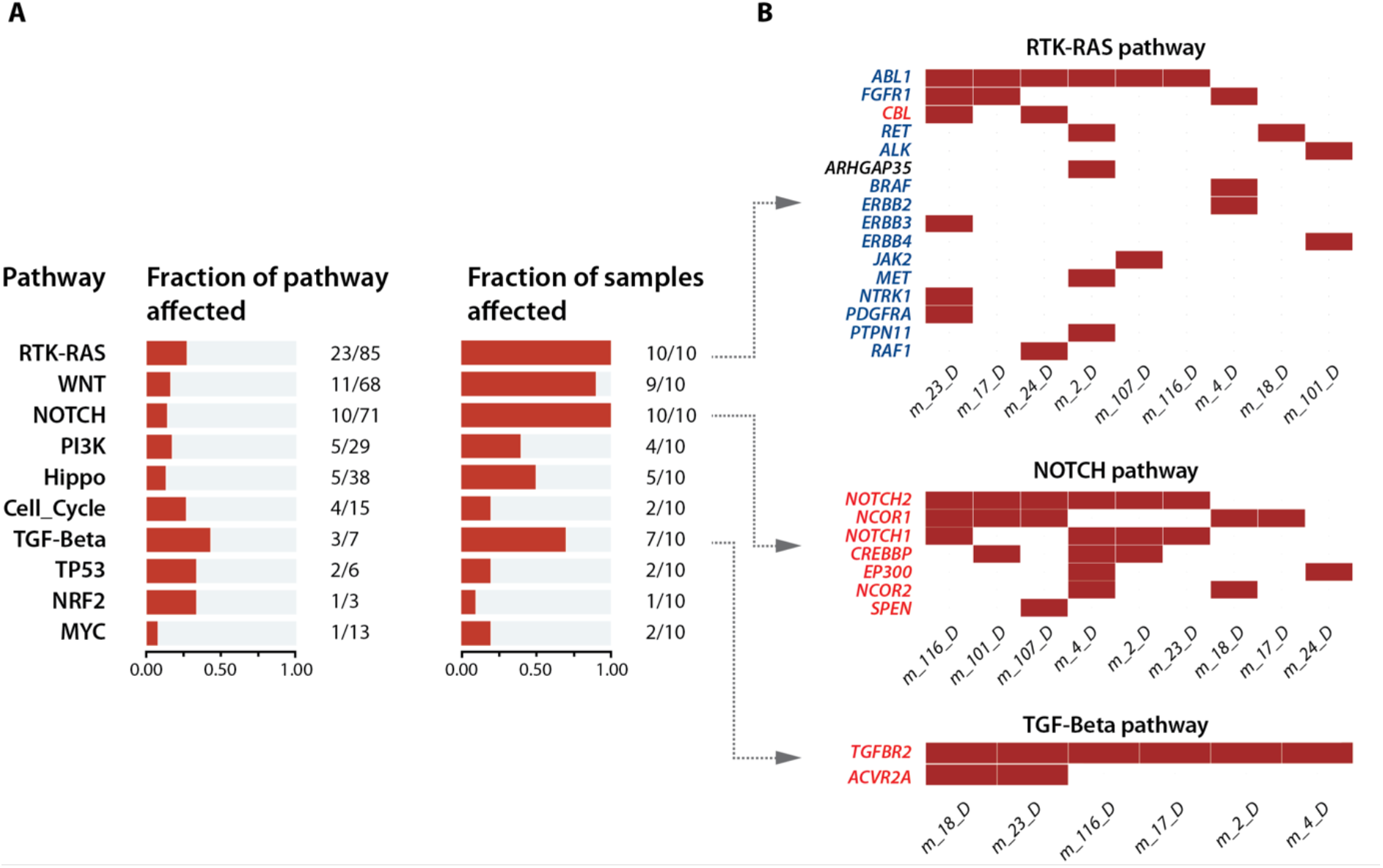
Overview of oncogenic pathway alterations in the patient cohort. The oncogenic pathways affected in the AEL patients are shown (A) with the number of affected genes in relation to the total number of genes linked to each pathway. The second plot displays the fraction of patients bearing mutations in a given pathway. (B) For each pathway, Var| Decrypt allows visualizing the mutated genes for each patient to easily spot the recurrently mutated genes. Oncogenes and tumor suppressor genes are indicated in blue and red, respectively.

### Visualization of mutational hotspots and amino acid changes

Finally, Var|Decrypt provides a visualization tool depicting the localization of mutations on a given gene product (protein). The known protein domains are displayed along with the position of the various mutations or variants detected, with a color code indicating the variant types (STOP gain, frameshifts, non-synonymous SNPs). This feature allows to detect mutational hotspots and preferential localization of mutations in functional protein domains, as shown in **Fig.7A** in the succinate dehydrogenase complex flavoprotein subunit A (*SDHA*) gene. In addition, a table provides the identity of amino acid changes along with several variant metrics (**Supp Table. 2**).

### Discovery of putative novel oncogenic mutations in AEL

We applied Var|Decrypt to decipher the mutational landscape of AEL. Besides the known and recently described mutations in *TET2* (40% of patients in our cohort), *TP53* (30% patients), *EZH2* (10%), *NCOR1/2* (50%/20%) or *GATA1* (in 10% of the patients), our tool highlighted mutational hotspots in several additional genes, likely representing important components of the AEL mutational landscape. We identified several mutations within the *SDHA* gene (**Fig. 7A**), a critical member of the succinate dehydrogenase complex, which were not primarily identified in the previous AEL studies (Iacobucci *et al*., 2019a; Fagnan *et al*., 2020; Cervera *et al*., 2016, 2017)(Grossmann *et al*., 2013). The succinate dehydrogenase (SDH) complex is a mitochondria-localized multiprotein complex involved in cellular respiration through the electron transfer chain (complex II) (Sharma *et al*., 2020, 2019). The SDH is nuclearly encoded and composed of 4 subunits (SDHA-D). Loss of function of any of SDH subunits may associate with neuroendocrine tumors or neurodegenerative disorders such as Leigh’s disease (Sharma *et al*., 2019). We identified a mutational hotspot within the *SDHA* gene in 70% of patients (**Fig. 7A**). Interestingly all detected missense mutations (V446A, A449V, A454T, S456L, R465Q, A466T, C467S) cluster around the key *SDHA* active site residue SDHA^R451^ (Sharma *et al*., 2020) (**Fig.7B**). Although the precise functional impact of such mutations is currently unknown, some or all may significantly alter SDHA active site spatial conformation and lead to (partial) insufficiency. Measuring complex II activity in AEL cells and its requirement for leukemia development is out of the scope of this study but represents an interesting lead to follow, as complex II alterations may be of importance for the development or maintenance of AEL.

**Figure 7.**
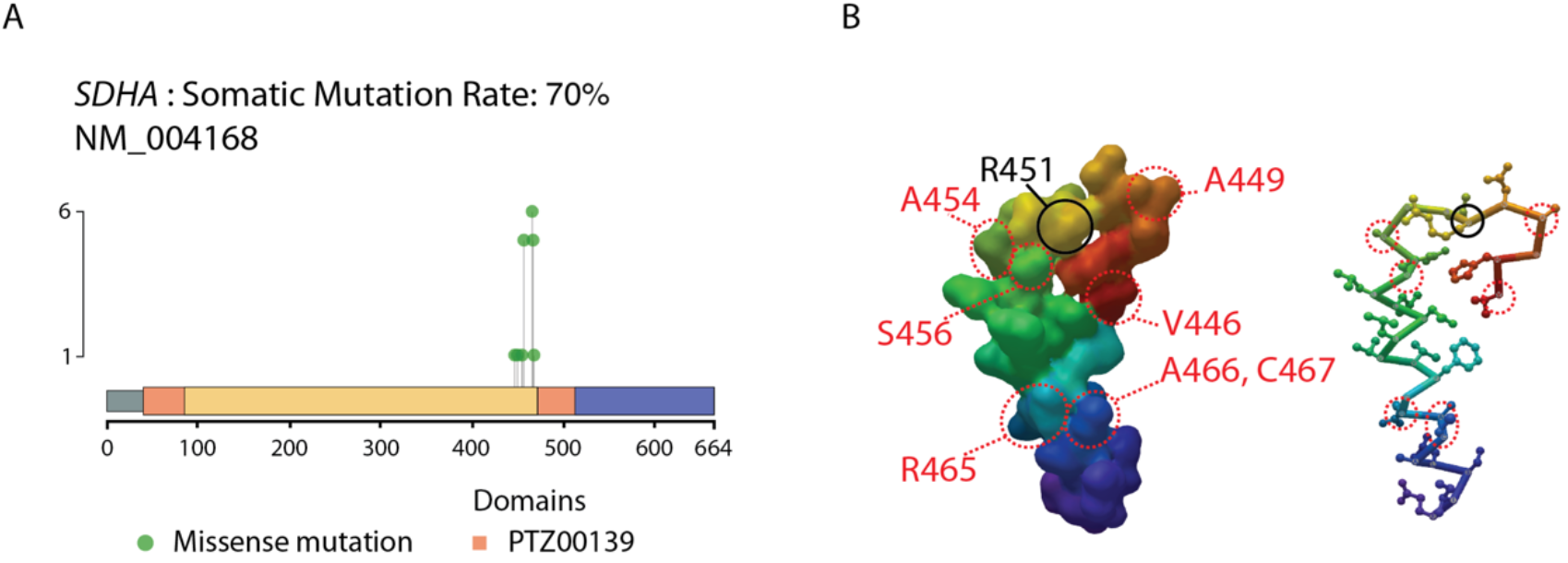
Visualizing mutational hotspots. (A) The 'amino-acid changes' page displays the protein domains with the localization and type of mutations in the entire cohort. The example of the SDHA gene is shown. (B) Structure (accession #6VAX) of the SDHA active site from (Sharma et al., 2020, 2). Only amino acids 446 to 472 are shown. The key active site residue R451 is indicated in black, the positions of the mutated residues in AEL are shown in red (left). Right, similar representation using the 'schematic view, with amino acid side chains shown as sticks and balls using MMDB viewer (Madej et al., 2014).

Finally, on a more global scale, Var|Decrypt allowed us to identify enrichment of mutations in oncogenic signaling pathways, in particular the receptor tyrosine kinase (RTK)/RAS pathway, with a high prevalence of IRS1 mutations (40% of patients, including in-frame deletions, non-synonymous SNVs and a STOP-gain), and mutations in *FGFR1*, *RET*, *JAK2* (30, 20, 10 % of patients, respectively), or BRAF (10%). Importantly, our tool highlighted the Notch pathway as frequently mutated in AEL with Notch1 and Notch 2 receptor variants found in 40% and 60% of the patients, respectively.

Taken altogether these data highlight putative novel oncogenic processes and pathways in AEL, and underscore the usefulness of Var|Decrypt to provide leads for functional explorations.

### Benchmarking Var|Decrypt on an independent dataset of 90 multiple myeloma samples

We sought to benchmark Var|Decrypt on an independent dataset. To this aim, we analyzed published WES data from 30 human multiple myeloma cell lines (HMCLs) and primary multiple myeloma (MM) from 59 patients (Vikova *et al*., 2019). Previous analysis of these data revealed a prevalent TP53 mutational landscape and altered MAPK pathways. Reanalyzing this dataset with Var|Decrypt after filtering-out the putative false positive hits (i.e. highly mutated gene families such as MUC genes, see above) using the filtering options (frequency less than 4 mutations by gene within 20% of the cohort), and after crossing the mutated gene list with cancer gene databases (COSMIC, as in (Vikova *et al*., 2019)) led to very similar identification of MM mutated hits, with frequent *TP53* (47%), *KRAS* (40%), *NRAS* (30%), *ATM* (33%) alterations, and many epigenetic modifiers (*BRD3*, *BRD4*, *SETD1B*) and DNA repair proteins (*FANCD2*, *RECQL4*) (**Supp. Table 3**). In particular we identify the MAPK/RAS pathway as recurrently altered (**Supp Fig. 2 and 3**) (Lohr *et al*., 2014; Bolli *et al*., 2014), validating the functionality of Var|Decrypt.

## Discussion

Although WES is widely used to diagnose human diseases or to discover pathological mutations in mendelian disorders and cancers, there is a surprising paucity of accessible and easy to use tools for WES analysis. A major hurdle is the presence of hundreds to thousands of variants routinely detected in WES datasets, complicating the identification of causative mutations and prioritization of variants in complex samples such as tumor biopsies. We present here Var|Decrypt, a fully automated dynamic interface for Exome-Seq data analysis. From a computational point of view, Var|Decrypt represents a fast, easy to use and flexible tool, which can be installed via Docker on several operating systems (Linux, macOS), downloaded (open-source scripts) to run via Rstudio, or directly used online. In addition, Var|Decrypt can be deployed on a Linux sever via the open-source Shiny Server software to be available to a large number of users. It is worth noting that the shiny server solution is suitable for institutions which need to centralize computing resources (Var|Decrypt docker image is configurated already to be deployed on a shiny sever). Thanks to its user-friendly interface and the variety of incorporated analysis options, Var|Decrypt can easily be used by both bioinformaticians and non-programming experts/wet lab scientists. It therefore fulfills a gap by providing a complete solution for WES data analyses.

The primary goal of our tool is to provide non specialists with a ready-to-use solution for easy browsing and exploration of Exome-Seq datasets. The rational is that driver mutations or gene mutations that are important for the development or maintenance of a disease state should be overrepresented or enriched in a patient cohort. It is still currently a major challenge to accurately and consistently classify variant pathogenicity (Richards *et al*., 2015; McInnes *et al*., 2021; Nicora *et al*., 2022). Although we do not directly address the pathogenicity of detected variants, we reasoned that focusing on the recurrently mutated genes may provide an increased likelihood of identifying important disease targets. Indeed, finding multiple different variants (possibly being of unknown pathogenicity) affecting the same gene product in the patient cohort would increase the chance to highlight disease drivers. In that respect, rather than focusing on the variants themselves, Var|Decrypt provides a gene centered analysis, allowing distinct variants affecting the same gene to score similarly. Hence the resulting tables and analyses provide users with a quick overview of the recurrently mutated genes and associated functions (pathways), representing putative disease drivers. The numerous built-in features allow extraction of such meaningful information e.g. recurrently mutated genes, or recurrently mutated pathway components, allowing to shed light on the mechanistic features of human disorders. We provide example of the analysis of AEL, a rare, poorly characterized and particularly aggressive subtype of leukemia. While recent work have started to shed light on the mutational landscape of AEL (Fagnan *et al*., 2020; Cervera *et al*., 2016, 2017; Grossmann *et al*., 2013; Iacobucci *et al*., 2019b; Micci *et al*., 2013), we show here that frequent NOTCH pathway alteration is associated with AEL in our cohort. We also identified mutational hotspots in the mitochondrial complex II component SDHA. This underscores the utility of dedicated analysis tools such as Var|Decrypt to highlight oncogenic signaling pathways and mutated genes that have been overlooked in other studies. Whether NOTCH pathway alteration represents a common feature of AEL and can be exploited as therapeutic vulnerability remains to be tested and is beyond the scope of this article. However, this represents an example of how biological information can be extracted from complex Exome-seq datasets without knowledge in bioinformatics. We further showed that Var|Decrypt could detect known recurrently mutated genes in human multiple myeloma, and could highlight signaling pathways known to be important for this disease. Taken altogether, these data indicate that Var|Decrypt represents a functional and attractive tool allowing efficient analysis of WES data, with the overarching goal to facilitate functional studies and guide therapeutic decisions. We expect that thanks to its ease of use and simple user-friendly interface, Var|Decrypt will help wet-lab scientists to get critical insight into the molecular mechanisms of human disorders.

## Acknowledgements

We wish to thank Samad El Kaoutari for assistance in the initial steps of Exome-Seq analysis pipeline development, and the GOELAMS for granting access to primary AEL samples.

## Funding

This work was supported by the Laboratory of excellence (Labex) EpiGenMed [program “Investissements d’avenir” ANR-10-LABX-12-01]; the Laboratory of Excellence GR-Ex [program “Investissements d’avenir” ANR-11-LABX-0051, ANR-18-IDEX-0001]; the Fondation pour la Recherche Médicale [Equipe FRM DEQ20180339221]; and the Ligue Nationale Contre le Cancer, ANR-18-CE15-0010-01 PLASMADIFF-3D, FFRMG (AAP-FFRMG-2021), INSERM PSCI 2020 Smooth-MM, and Institut Universitaire de France.

## Key Points

- Var|Decrypt offers an all-in-one solution tool for WES functional analysis
- Var|Decrypt aims at bridging the gap between computational scientists and wet lab scientists & clinicians
- Easy to use and user-friendly interface incorporating a multitude of functional enrichment tools for the discovery of disease drivers

## Methods

### Data

We used WES data from primary erythroleukemia samples (Fagnan *et al*., 2020). The data are separated into two types, namely ‘Tumoral’ samples, being the patient leukemic blasts samples, and ‘Normal’ representing the non-leukemic matched controls, considered healthy (non-leukemic marrow cells), and used for variant filtering purposes. So for each patient, matched leukemic sample (tumoral) and a control sample (normal) are used.

### Whole Exome Sequencing analysis pipeline

The human reference genome hg19 (https://hgdownload.cse.ucsc.edu/goldenPath/hg19/bigZips/) was indexed by BWA. Reads were trimmed with Trimmomatic (version0.36) to eliminate sequencing adapters and low-quality reads. Mapping was performed using BWA-MEM (version 0.7.17) with default parameters. SAM files were converted, sorted and indexed by Samtools (version 1.9). To improve alignment quality, MarkDuplicates tool from PICARD (version 2.17.11) was used to locate and tag duplicate reads within the BAM. Before preforming base recalibration for BAM files by BaseRecalibrator from GATK (version v3.8-1-0), reads were processed by AddOrReplaceReadGroups from PICARD (version 2.17.11) to define a group for all reads generated from the same run. For the variant calling step, the HaplotypeCaller was used for germline variants and Mutect2 for somatic ones from GATK4 (version 4.0.3.0) (Poplin *et al*., 2018). Variant filtering and annotation were performed by ANNOVAR (version Sun, 7 Jun 2020). First, results from the previous step are filtered by extracting all the known SNV involved in one or more human diseases (regardless of disease using the option: – filter -dbtype clinvar_20200316). Then, all known human variants were ignored (-filter -dbtype 1000g2015aug_all) (Wang *et al*., 2010). Finally, all unknown variants were grouped with the pathogenic variants for each patient/sample data. After the annotation step by ANNOVAR, only exonic polymorphisms were considered and kept in the output file. Using custom Python scripts, synonymous variants were removed from the final output files. The resulting files were used as input files for Var|Decrypt.

### Exome Sequencing analyses using Var|Decrypt

Downstream analyses and variants prioritization were performed using Var|Decrypt, which mainly uses the following R libraries:

(1) Shiny, which allows to develop a user-friendly graphical interface to visualize the various types of data. This application can be launched in Rstudio or in any modern web browser such as firefox, chrome or safari (Chang *et al*., 2020); (2) DOSE which allows to perform enrichment analyses of a set of genes to discover gene-disease associations. It implements several methods to measure the semantic similarities between the DO (Disease ontology) terms and the different gene products (Yu *et al*., 2015); (3) clusterProfiler, which implements methods for analyzing and visualizing functional profiles from gene clusters (Yu *et al*., 2012); (4) org.Hs.eg.db, which contains annotation of the human genome (Carlson, 2019); (5) ReactomePA, which provides signaling pathway analysis functions based on the REACTOME database with several visualization functions (Yu and He, 2016); (6) networkD3, which was used to generate reactive 3D networks (Allaire *et al*., 2017). Finally, Var|Decrypt also uses maftools R package to process somatic variants (Mayakonda *et al*., 2018).

## Notes

### Competing Interest Statement

The authors have declared no competing interest.

