## Supplementary Data file for "Var|Decrypt: a novel and user-friendly tool to explore and prioritize variants in whole-exome sequencing data"

* correspondance :

MS :
ES :

### Supplementary Figures and Tables

**Supplementary Figure 1.** Example of Var|Decrypt front page showing the mutated gene list together with the frequencies in the cohort and the mutation types.

**Supplementary Figure 2.** Analysis of WES from 30 human multiple myeloma cell lines. (A) Altered pathways overrepresented in the HMCL mutated genes. (B) mutations in oncogenic signaling pathways showing that the RTK-RAS and NOTCH pathways are among the top mutated pathways.

**Supplementary Figure 3.** Detailed view and frequencies of the RTK-RAS pathway mutated genes. Affected genes belonging to the RTK-RAS pathway are shown and highlighted when mutated for each sample. (A) The figure shows prevalent FGFR4 (17 out of 29 samples, 58%), and KRAS (12 out of 29 samples, 41%) mutations in HMCL, and (B) similar frequencies (FGFR4 38/59, 64%; KRAS 17/59, 28%) were observed in primary human multiple myeloma samples.

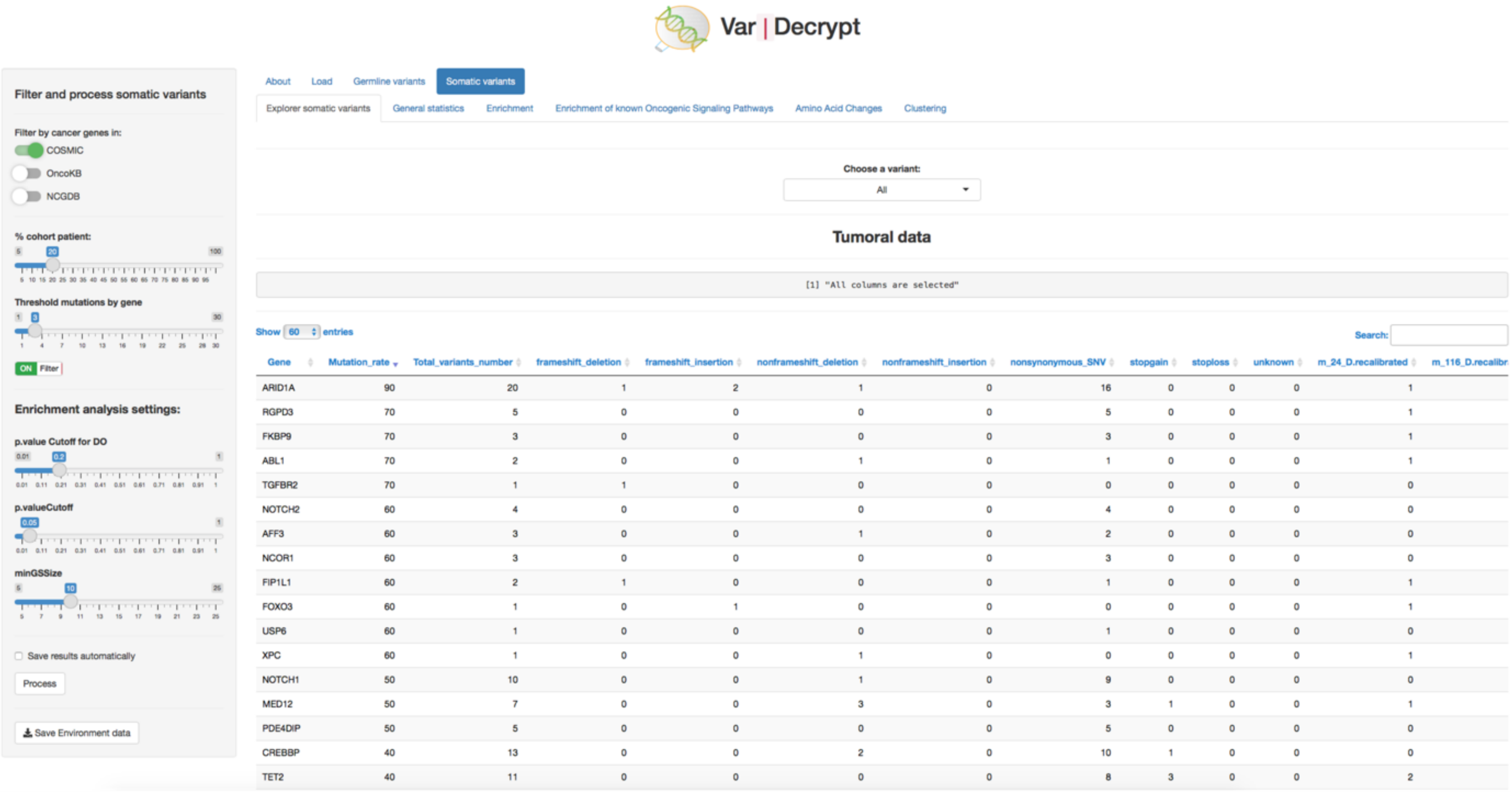


Supplementary Figure 1. Example of Var|Decrypt front page showing the mutated gene list together with the frequencies in the cohort and the mutation types.

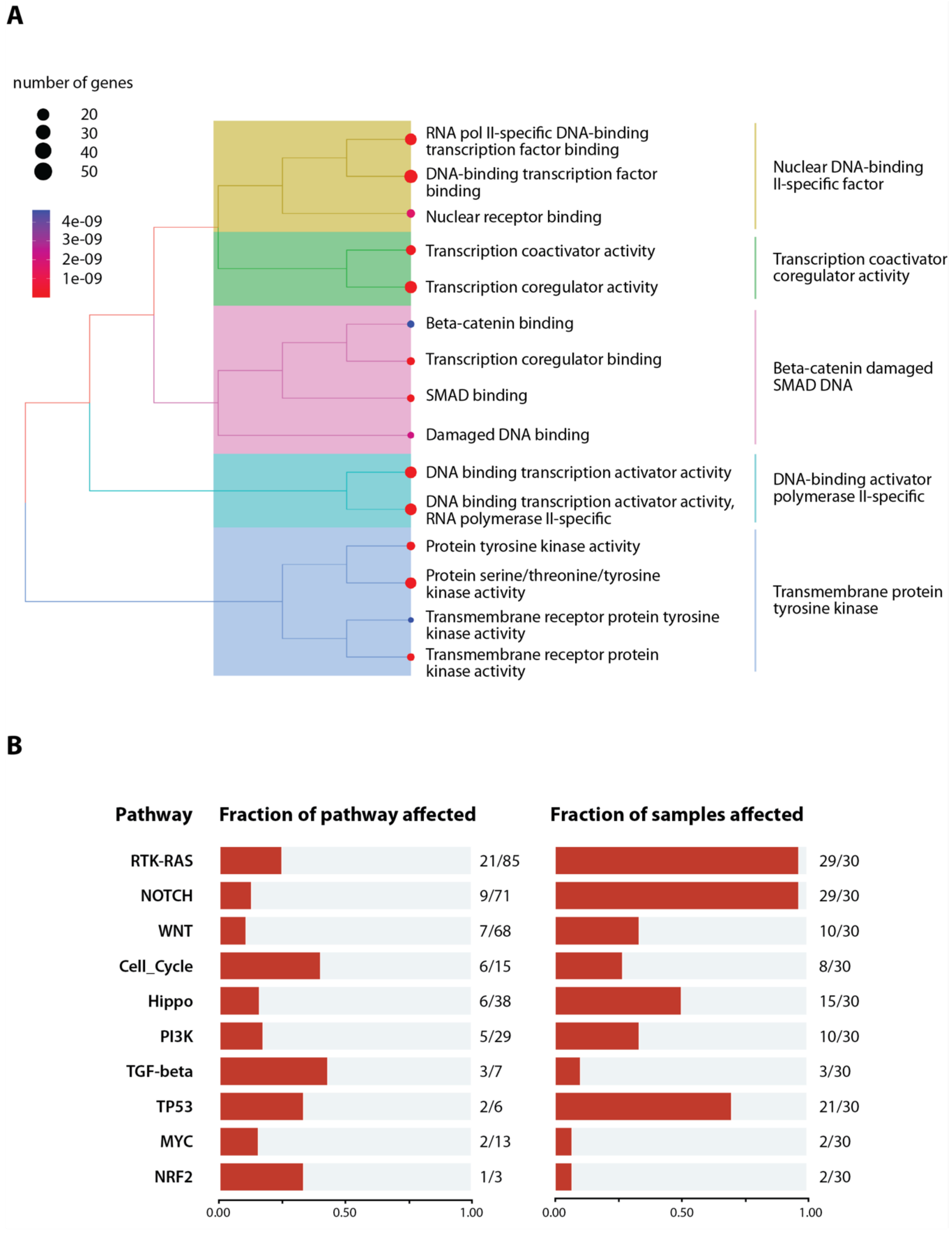


Supplementary Figure 2. Analysis of WES from 30 human multiple myeloma cell lines. (A) Altered pathways overrepresented in the HMCL mutated genes. (B) mutations in oncogenic signaling pathways showing that the RTK-RAS and NOTCH pathways are among the top mutated pathways.

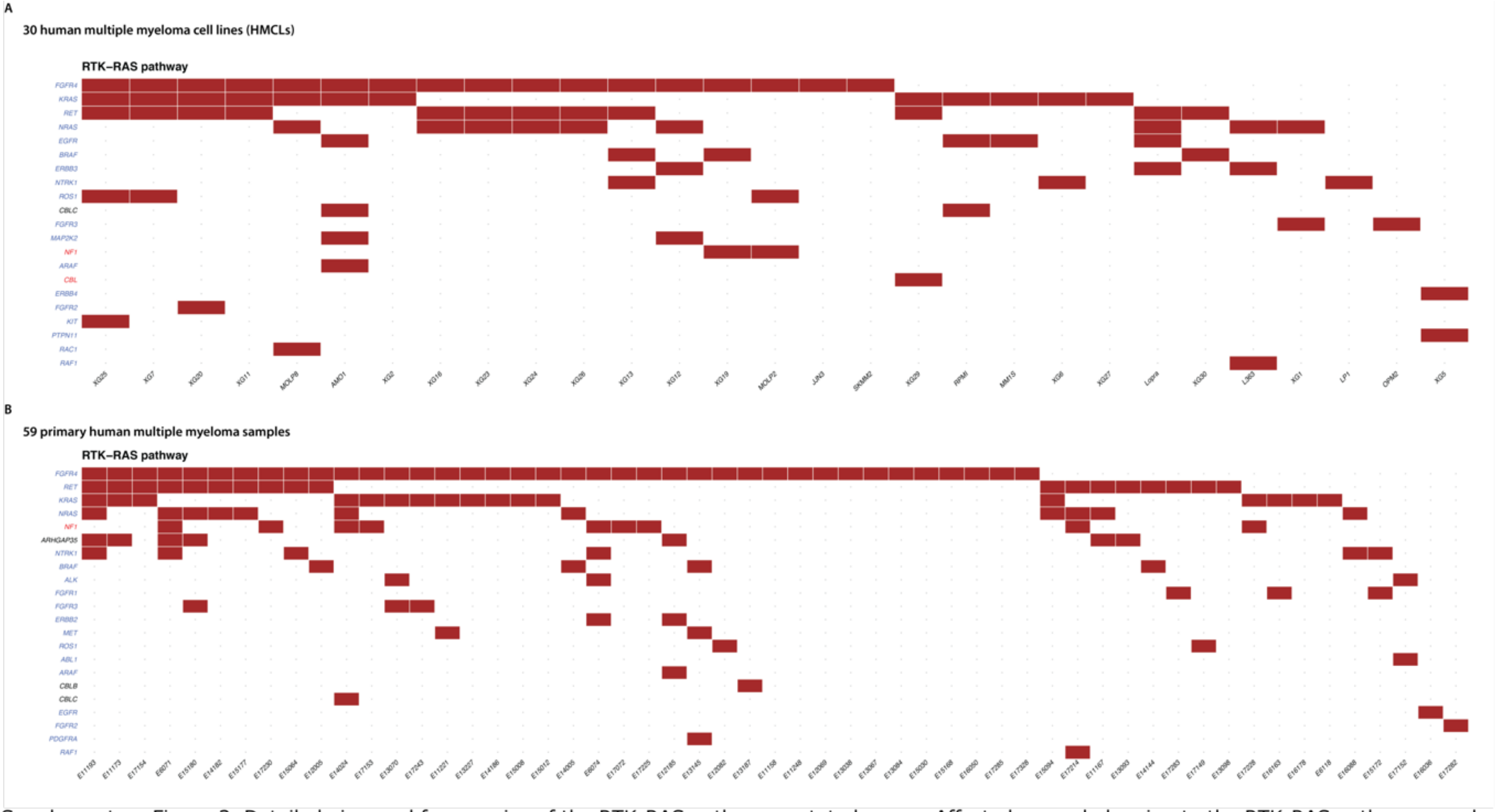


Supplementary Figure 3. Detailed view and frequencies of the RTK-RAS pathway mutated genes. Affected genes belonging to the RTK-RAS pathway are shown and highlighted when mutated for each sample. (A) The figure shows prevalent FGFR4 (17 out of 29 samples, 58%), and KRAS (12 out of 29 samples, 41%) mutations in HMCL, and (B) similar frequencies (FGFR4 38/59, 64%; KRAS 17/59, 28%) were observed in primary human multiple myeloma samples.

**Supplementary Table 1. Top 100 mutated genes in the AEL cohort.**
The table depicts the Var I Decrypt output with the top 100 mutated genes in AEL patients, with detailed information of the mutation types such as frameshifts, stop-gains, etc.gry columns ) and the number of mutations in each sample/patient (yellow columns).

| | | | VARIANT TYPES | | | | | | | | PATIENT ID | | | | | | | | | |
|---|---|---|---|---|---|---|---|---|---|---|---|---|---|---|---|---|---|---|---|---|
| Gene | Mutation rate (%) | Total number of variants | Frameshift deletion | Frameshift insertion | Non-frameshift deletion | Non-frameshift insertion | Non-synonymous SNV | STOP gain | STOP loss | unknown | m_24_D | m_116_D | m_4_D | m_23_D | m_18_D | m_13_D | m_107_D | m_101_D | m_17_D | m_2_D |
| ARID1A | 90 | 20 | 1 | 2 | 1 | 0 | 16 | 0 | 0 | 0 | 1 | 1 | 1 | 1 | 1 | 20 | 0 | 1 | 1 | 1 |
| RGPD3 | 70 | 5 | 0 | 0 | 0 | 0 | 5 | 0 | 0 | 0 | 1 | 2 | 0 | 1 | 1 | 2 | 3 | 1 | 0 | 0 |
| FKBP9 | 70 | 3 | 0 | 0 | 0 | 0 | 3 | 0 | 0 | 0 | 1 | 0 | 1 | 0 | 1 | 1 | 1 | 1 | 0 | 2 |
| ABL1 | 70 | 2 | 0 | 0 | 1 | 0 | 1 | 0 | 0 | 0 | 1 | 2 | 0 | 1 | 0 | 1 | 1 | 0 | 1 | 1 |
| TGFBR2 | 70 | 1 | 1 | 0 | 0 | 0 | 0 | 0 | 0 | 0 | 0 | 1 | 1 | 1 | 1 | 1 | 0 | 0 | 1 | 1 |
| NOTCH2 | 60 | 4 | 0 | 0 | 0 | 0 | 4 | 0 | 0 | 0 | 0 | 1 | 1 | 1 | 0 | 0 | 1 | 2 | 0 | 1 |
| AFF3 | 60 | 3 | 0 | 0 | 1 | 0 | 2 | 0 | 0 | 0 | 0 | 1 | 0 | 0 | 1 | 2 | 1 | 0 | 1 | 1 |
| NCOR1 | 60 | 3 | 0 | 0 | 0 | 0 | 3 | 0 | 0 | 0 | 0 | 1 | 0 | 0 | 1 | 1 | 1 | 2 | 1 | 0 |
| FIP1L1 | 60 | 2 | 1 | 0 | 0 | 0 | 1 | 0 | 0 | 0 | 1 | 1 | 0 | 1 | 1 | 2 | 0 | 0 | 0 | 1 |
| FOXO3 | 60 | 1 | 0 | 1 | 0 | 0 | 0 | 0 | 0 | 0 | 1 | 1 | 0 | 0 | 1 | 1 | 1 | 0 | 1 | 0 |
| USP6 | 60 | 1 | 0 | 0 | 0 | 0 | 1 | 0 | 0 | 0 | 0 | 1 | 1 | 1 | 1 | 1 | 0 | 0 | 0 | 1 |
| XPC | 60 | 1 | 0 | 0 | 1 | 0 | 0 | 0 | 0 | 0 | 1 | 1 | 1 | 0 | 1 | 0 | 1 | 1 | 0 | 0 |
| NOTCH1 | 50 | 10 | 0 | 0 | 1 | 0 | 9 | 0 | 0 | 0 | 0 | 1 | 2 | 1 | 0 | 6 | 0 | 0 | 0 | 1 |
| MED12 | 50 | 7 | 0 | 0 | 3 | 0 | 3 | 1 | 0 | 0 | 1 | 1 | 0 | 0 | 2 | 3 | 0 | 0 | 0 | 1 |
| PDE4DIP | 50 | 5 | 0 | 0 | 0 | 0 | 5 | 0 | 0 | 0 | 0 | 1 | 0 | 1 | 0 | 3 | 1 | 1 | 0 | 0 |
| CREBBP | 40 | 13 | 0 | 0 | 2 | 0 | 10 | 1 | 0 | 0 | 0 | 0 | 1 | 0 | 0 | 10 | 0 | 1 | 0 | 1 |
| TET2 | 40 | 11 | 0 | 0 | 0 | 0 | 8 | 3 | 0 | 0 | 2 | 0 | 1 | 0 | 0 | 6 | 0 | 0 | 2 | 0 |
| CACNA1D | 40 | 9 | 1 | 0 | 0 | 0 | 8 | 0 | 0 | 0 | 0 | 0 | 0 | 0 | 1 | 6 | 0 | 0 | 1 | 2 |
| MN1 | 40 | 4 | 0 | 0 | 3 | 0 | 1 | 0 | 0 | 0 | 0 | 0 | 3 | 0 | 2 | 1 | 1 | 0 | 0 | 0 |
| CTNND2 | 40 | 4 | 0 | 0 | 1 | 0 | 3 | 0 | 0 | 0 | 0 | 0 | 0 | 1 | 1 | 3 | 0 | 1 | 0 | 0 |
| DNM2 | 40 | 2 | 0 | 0 | 0 | 0 | 2 | 0 | 0 | 0 | 1 | 0 | 2 | 1 | 0 | 1 | 0 | 0 | 0 | 0 |
| CHD4 | 40 | 2 | 0 | 0 | 1 | 0 | 1 | 0 | 0 | 0 | 1 | 0 | 1 | 2 | 1 | 0 | 0 | 0 | 0 | 0 |
| ZNRF3 | 40 | 1 | 0 | 0 | 1 | 0 | 0 | 0 | 0 | 0 | 0 | 1 | 0 | 0 | 1 | 0 | 1 | 1 | 0 | 0 |
| MACC1 | 40 | 1 | 0 | 0 | 0 | 0 | 1 | 0 | 0 | 0 | 1 | 0 | 1 | 1 | 0 | 1 | 0 | 0 | 0 | 0 |
| KAT6B | 30 | 27 | 3 | 0 | 2 | 0 | 21 | 1 | 0 | 0 | 0 | 0 | 1 | 1 | 0 | 25 | 0 | 0 | 0 | 0 |
| BCOR | 30 | 26 | 2 | 2 | 0 | 0 | 22 | 0 | 0 | 0 | 0 | 0 | 0 | 1 | 0 | 24 | 0 | 0 | 0 | 1 |
| BCORL1 | 30 | 25 | 0 | 0 | 0 | 1 | 24 | 0 | 0 | 0 | 0 | 1 | 1 | 0 | 0 | 23 | 0 | 0 | 0 | 0 |
| KMT2D | 30 | 14 | 1 | 1 | 1 | 0 | 11 | 0 | 0 | 0 | 0 | 0 | 0 | 0 | 0 | 13 | 1 | 0 | 0 | 1 |
| BCL11A | 30 | 8 | 0 | 0 | 0 | 0 | 8 | 0 | 0 | 0 | 0 | 0 | 0 | 0 | 0 | 6 | 0 | 1 | 0 | 2 |
| EP300 | 30 | 7 | 0 | 0 | 0 | 1 | 6 | 0 | 0 | 0 | 1 | 0 | 1 | 0 | 0 | 5 | 0 | 0 | 0 | 0 |
| CBL | 30 | 6 | 0 | 0 | 2 | 0 | 4 | 0 | 0 | 0 | 1 | 0 | 0 | 1 | 0 | 5 | 0 | 0 | 0 | 0 |
| ARHGEF12 | 30 | 6 | 0 | 0 | 1 | 1 | 4 | 0 | 0 | 0 | 0 | 0 | 0 | 0 | 0 | 4 | 1 | 0 | 0 | 1 |
| ATR | 30 | 5 | 1 | 0 | 0 | 0 | 3 | 1 | 0 | 0 | 0 | 1 | 0 | 0 | 0 | 3 | 0 | 1 | 0 | 0 |
| ATP2B3 | 30 | 4 | 0 | 0 | 0 | 0 | 4 | 0 | 0 | 0 | 0 | 0 | 1 | 0 | 0 | 2 | 0 | 0 | 0 | 1 |
| IRF4 | 30 | 4 | 0 | 0 | 0 | 0 | 4 | 0 | 0 | 0 | 0 | 0 | 1 | 0 | 0 | 2 | 0 | 0 | 0 | 3 |
| PBRM1 | 30 | 4 | 0 | 0 | 0 | 0 | 4 | 0 | 0 | 0 | 0 | 0 | 1 | 0 | 0 | 2 | 1 | 0 | 0 | 0 |
| NCOR2 | 30 | 3 | 0 | 0 | 0 | 0 | 3 | 0 | 0 | 0 | 0 | 0 | 1 | 0 | 1 | 1 | 0 | 0 | 0 | 0 |
| BAZ1A | 30 | 3 | 0 | 0 | 0 | 0 | 2 | 1 | 0 | 0 | 0 | 0 | 1 | 0 | 1 | 1 | 0 | 0 | 0 | 0 |
| FGFR1 | 30 | 2 | 0 | 0 | 1 | 0 | 1 | 0 | 0 | 0 | 0 | 0 | 1 | 1 | 0 | 0 | 0 | 0 | 1 | 0 |
| HSP90AA1 | 30 | 1 | 0 | 0 | 1 | 0 | 0 | 0 | 0 | 0 | 0 | 0 | 1 | 0 | 0 | 0 | 0 | 1 | 1 | 0 |
| ACVR2A | 30 | 1 | 0 | 0 | 0 | 0 | 1 | 0 | 0 | 0 | 0 | 0 | 0 | 1 | 1 | 1 | 0 | 0 | 0 | 0 |
| CTNNA2 | 30 | 1 | 0 | 0 | 0 | 0 | 1 | 0 | 0 | 0 | 0 | 1 | 0 | 0 | 0 | 0 | 1 | 1 | 0 | 0 |
| TSC1 | 30 | 1 | 0 | 0 | 1 | 0 | 0 | 0 | 0 | 0 | 0 | 0 | 0 | 0 | 0 | 0 | 1 | 1 | 0 | 1 |
| ASXL1 | 20 | 18 | 2 | 2 | 0 | 0 | 13 | 1 | 0 | 0 | 1 | 0 | 0 | 0 | 0 | 17 | 0 | 0 | 0 | 0 |
| USP9X | 20 | 17 | 1 | 1 | 0 | 0 | 15 | 0 | 0 | 0 | 0 | 0 | 1 | 0 | 0 | 16 | 0 | 0 | 0 | 0 |
| THRAP3 | 20 | 12 | 1 | 1 | 0 | 0 | 10 | 0 | 0 | 0 | 0 | 0 | 1 | 0 | 0 | 11 | 0 | 0 | 0 | 0 |
| KMT2A | 20 | 12 | 0 | 0 | 0 | 0 | 12 | 0 | 0 | 0 | 0 | 0 | 1 | 0 | 0 | 11 | 0 | 0 | 0 | 0 |
| BCL9 | 20 | 10 | 0 | 0 | 0 | 0 | 10 | 0 | 0 | 0 | 0 | 0 | 0 | 1 | 0 | 10 | 0 | 0 | 0 | 0 |
| GRIN2A | 20 | 10 | 0 | 0 | 0 | 0 | 10 | 0 | 0 | 0 | 0 | 0 | 0 | 0 | 0 | 9 | 0 | 0 | 1 | 0 |
| ARID2 | 20 | 8 | 0 | 0 | 0 | 0 | 8 | 0 | 0 | 0 | 0 | 0 | 0 | 0 | 0 | 7 | 0 | 0 | 0 | 1 |
| RBM10 | 20 | 8 | 0 | 0 | 1 | 0 | 6 | 1 | 0 | 0 | 0 | 1 | 0 | 0 | 0 | 7 | 0 | 0 | 0 | 0 |
| IRS4 | 20 | 8 | 0 | 0 | 0 | 0 | 8 | 0 | 0 | 0 | 0 | 0 | 1 | 0 | 0 | 7 | 0 | 0 | 0 | 0 |
| ETV6 | 20 | 8 | 0 | 3 | 0 | 0 | 5 | 0 | 0 | 0 | 0 | 0 | 1 | 0 | 0 | 7 | 0 | 0 | 0 | 0 |
| ABL2 | 20 | 8 | 0 | 0 | 0 | 0 | 8 | 0 | 0 | 0 | 0 | 0 | 2 | 0 | 0 | 6 | 0 | 0 | 0 | 0 |
| ASXL2 | 20 | 7 | 0 | 0 | 1 | 0 | 6 | 0 | 0 | 0 | 0 | 0 | 0 | 0 | 0 | 6 | 0 | 0 | 0 | 1 |
| DDX3X | 20 | 7 | 0 | 0 | 0 | 0 | 7 | 0 | 0 | 0 | 0 | 0 | 0 | 0 | 0 | 6 | 0 | 0 | 0 | 1 |
| POLQ | 20 | 7 | 1 | 1 | 0 | 0 | 5 | 0 | 0 | 0 | 0 | 0 | 0 | 0 | 0 | 1 | 0 | 0 | 0 | 6 |
| BCL11B | 20 | 7 | 0 | 0 | 1 | 1 | 5 | 0 | 0 | 0 | 0 | 1 | 0 | 0 | 0 | 6 | 0 | 0 | 0 | 0 |
| PRDM16 | 20 | 7 | 0 | 0 | 0 | 0 | 7 | 0 | 0 | 0 | 0 | 0 | 1 | 0 | 0 | 6 | 0 | 0 | 0 | 0 |
| PRCC | 20 | 7 | 0 | 0 | 0 | 0 | 7 | 0 | 0 | 0 | 0 | 0 | 0 | 0 | 0 | 7 | 0 | 0 | 0 | 1 |
| ATRX | 20 | 7 | 0 | 0 | 0 | 0 | 7 | 0 | 0 | 0 | 0 | 0 | 0 | 0 | 0 | 6 | 0 | 0 | 0 | 1 |
| TFE3 | 20 | 7 | 0 | 0 | 0 | 0 | 7 | 0 | 0 | 0 | 0 | 0 | 1 | 0 | 0 | 6 | 0 | 0 | 0 | 0 |
| BRAF | 20 | 6 | 1 | 1 | 0 | 0 | 4 | 0 | 0 | 0 | 0 | 0 | 1 | 0 | 0 | 5 | 0 | 0 | 0 | 0 |
| SOX2 | 20 | 6 | 0 | 0 | 0 | 1 | 5 | 0 | 0 | 0 | 0 | 0 | 0 | 0 | 0 | 5 | 0 | 1 | 0 | 0 |
| TRIM33 | 20 | 6 | 0 | 0 | 0 | 0 | 6 | 0 | 0 | 0 | 0 | 0 | 0 | 0 | 0 | 6 | 0 | 0 | 0 | 2 |
| TCF12 | 20 | 5 | 0 | 0 | 0 | 0 | 5 | 0 | 0 | 0 | 0 | 0 | 0 | 0 | 0 | 4 | 0 | 0 | 0 | 1 |
| CSMD3 | 20 | 5 | 0 | 0 | 0 | 0 | 5 | 0 | 0 | 0 | 0 | 0 | 0 | 0 | 0 | 4 | 0 | 0 | 0 | 1 |
| MYB | 20 | 5 | 0 | 0 | 0 | 0 | 5 | 0 | 0 | 0 | 0 | 0 | 0 | 0 | 0 | 4 | 0 | 1 | 0 | 0 |
| DICER1 | 20 | 5 | 0 | 0 | 0 | 0 | 4 | 1 | 0 | 0 | 0 | 0 | 0 | 0 | 0 | 4 | 0 | 0 | 0 | 1 |
| SPEN | 20 | 4 | 0 | 0 | 0 | 0 | 4 | 0 | 0 | 0 | 0 | 0 | 0 | 0 | 0 | 3 | 1 | 0 | 0 | 0 |
| AFF1 | 20 | 4 | 2 | 1 | 0 | 0 | 1 | 0 | 0 | 0 | 1 | 0 | 0 | 0 | 0 | 0 | 0 | 0 | 0 | 3 |
| EZH2 | 20 | 4 | 0 | 1 | 0 | 0 | 3 | 0 | 0 | 0 | 1 | 0 | 0 | 0 | 0 | 3 | 0 | 0 | 0 | 0 |
| BMP5 | 20 | 4 | 1 | 0 | 0 | 0 | 3 | 0 | 0 | 0 | 0 | 0 | 0 | 0 | 0 | 3 | 0 | 0 | 0 | 1 |
| PRDM2 | 20 | 4 | 0 | 0 | 0 | 0 | 4 | 0 | 0 | 0 | 0 | 0 | 1 | 0 | 0 | 3 | 0 | 0 | 0 | 0 |
| MYC | 20 | 4 | 0 | 0 | 1 | 0 | 3 | 0 | 0 | 0 | 0 | 1 | 0 | 0 | 0 | 3 | 0 | 0 | 0 | 0 |
| CIC | 20 | 4 | 0 | 0 | 0 | 0 | 4 | 0 | 0 | 0 | 0 | 0 | 0 | 0 | 0 | 3 | 0 | 0 | 1 | 0 |
| ARHGAP35 | 20 | 4 | 0 | 0 | 1 | 0 | 3 | 0 | 0 | 0 | 0 | 0 | 0 | 0 | 0 | 4 | 0 | 0 | 0 | 1 |
| LCP1 | 20 | 3 | 0 | 0 | 0 | 0 | 3 | 0 | 0 | 0 | 0 | 0 | 1 | 0 | 0 | 2 | 0 | 0 | 0 | 0 |
| SKI | 20 | 3 | 0 | 0 | 0 | 0 | 3 | 0 | 0 | 0 | 0 | 0 | 0 | 0 | 0 | 2 | 0 | 0 | 0 | 1 |
| BCLAF1 | 20 | 3 | 0 | 0 | 0 | 0 | 2 | 1 | 0 | 0 | 0 | 1 | 0 | 0 | 0 | 2 | 0 | 0 | 0 | 0 |
| CNTNAP2 | 20 | 3 | 0 | 0 | 1 | 0 | 2 | 0 | 0 | 0 | 0 | 0 | 0 | 0 | 0 | 2 | 0 | 0 | 0 | 1 |
| BCR | 20 | 3 | 0 | 1 | 0 | 0 | 2 | 0 | 0 | 0 | 0 | 0 | 0 | 0 | 0 | 2 | 1 | 0 | 0 | 0 |
| DDR2 | 20 | 3 | 0 | 0 | 0 | 0 | 3 | 0 | 0 | 0 | 0 | 0 | 1 | 0 | 0 | 2 | 0 | 0 | 0 | 0 |
| KIAA1549 | 20 | 3 | 1 | 1 | 0 | 0 | 1 | 0 | 0 | 0 | 1 | 0 | 0 | 0 | 0 | 2 | 0 | 0 | 0 | 0 |
| TPR | 20 | 3 | 0 | 0 | 0 | 0 | 2 | 1 | 0 | 0 | 0 | 0 | 0 | 0 | 0 | 2 | 0 | 0 | 0 | 1 |
| RAD21 | 20 | 3 | 0 | 1 | 0 | 0 | 2 | 0 | 0 | 0 | 0 | 0 | 0 | 0 | 0 | 2 | 0 | 0 | 1 | 0 |
| MLH1 | 20 | 3 | 0 | 0 | 0 | 0 | 3 | 0 | 0 | 0 | 0 | 0 | 0 | 0 | 0 | 2 | 0 | 1 | 0 | 0 |
| SETD2 | 20 | 3 | 0 | 0 | 0 | 0 | 3 | 0 | 0 | 0 | 0 | 0 | 0 | 0 | 0 | 2 | 0 | 0 | 0 | 1 |
| AFDN | 20 | 3 | 0 | 0 | 1 | 0 | 2 | 0 | 0 | 0 | 0 | 0 | 0 | 1 | 0 | 2 | 0 | 0 | 0 | 0 |
| PDGFRA | 20 | 2 | 0 | 0 | 0 | 0 | 2 | 0 | 0 | 0 | 0 | 0 | 0 | 1 | 0 | 1 | 0 | 0 | 0 | 0 |
| NIN | 20 | 2 | 0 | 0 | 0 | 0 | 1 | 1 | 0 | 0 | 0 | 0 | 1 | 0 | 0 | 0 | 0 | 0 | 0 | 1 |
| RAF1 | 20 | 2 | 0 | 0 | 0 | 0 | 2 | 0 | 0 | 0 | 1 | 0 | 0 | 0 | 0 | 1 | 0 | 0 | 0 | 0 |
| FBXO11 | 20 | 2 | 0 | 0 | 1 | 0 | 1 | 0 | 0 | 0 | 0 | 1 | 0 | 0 | 0 | 1 | 0 | 0 | 0 | 0 |
| ERBB3 | 20 | 2 | 0 | 0 | 0 | 0 | 2 | 0 | 0 | 0 | 0 | 0 | 0 | 1 | 0 | 1 | 0 | 0 | 0 | 0 |
| MAP2K4 | 20 | 2 | 0 | 0 | 0 | 0 | 2 | 0 | 0 | 0 | 0 | 0 | 0 | 0 | 0 | 1 | 0 | 0 | 0 | 1 |
| GNAQ | 20 | 2 | 0 | 0 | 0 | 0 | 1 | 1 | 0 | 0 | 0 | 0 | 0 | 0 | 0 | 0 | 0 | 0 | 1 | 2 |
| ROBO2 | 20 | 2 | 0 | 0 | 0 | 0 | 2 | 0 | 0 | 0 | 0 | 0 | 0 | 1 | 0 | 0 | 0 | 0 | 0 | 1 |
| RSPO3 | 20 | 2 | 0 | 0 | 0 | 0 | 2 | 0 | 0 | 0 | 0 | 0 | 0 | 0 | 0 | 1 | 0 | 0 | 1 | 0 |
| ARHGAP5 | 20 | 2 | 0 | 0 | 0 | 0 | 2 | 0 | 0 | 0 | 1 | 0 | 0 | 0 | 0 | 1 | 0 | 0 | 0 | 0 |
| ERBB4 | 20 | 2 | 0 | 0 | 0 | 0 | 2 | 0 | 0 | 0 | 0 | 0 | 0 | 0 | 0 | 1 | 0 | 1 | 0 | 0 |

**Supplementary Table 2. Variants and protein metrics.**

The table depicts the Var I Decrypt output for the protein and variat metrics indictiong the position of the mutations within the protein and cDNA sequence.

| Chromosome | Start_Position | End_Position | Reference Allele | Tumor Seq_Allele2 | Tumor Sample Barcode | Hugo Symbol | Variant Classification | tx | exon | txChange | aaChange | Variant_Type | Func.refGene | Gene.refGene | ExonicFunc.refGene | Amino acid change |
|---|---|---|---|---|---|---|---|---|---|---|---|---|---|---|---|---|
| chr4 | 106162520 | 106162520 | G | C | m_13_D | TET2 | Missense_Mutation | NM_001127208 | exon4 | c.G3434C | p.G1145A | SNP | exonic | TET2 | nonsynonymous SNV | TET2:NM_001127208:exon4:c.G3434C:p.G1145A |
| chr4 | 106162526 | 106162526 | T | A | m_13_D | TET2 | Missense_Mutation | NM_001127208 | exon4 | c.T3440A | p.F1147Y | SNP | exonic | TET2 | nonsynonymous SNV | TET2:NM_001127208:exon4:c.T3440A:p.F1147Y |
| chr4 | 106162552 | 106162552 | A | G | m_13_D | TET2 | Missense_Mutation | NM_001127208 | exon4 | c.A3466G | p.N1156D | SNP | exonic | TET2 | nonsynonymous SNV | TET2:NM_001127208:exon4:c.A3466G:p.N1156D |
| chr4 | 106162570 | 106162570 | G | A | m_13_D | TET2 | Missense_Mutation | NM_001127208 | exon4 | c.G3484A | p.E1162K | SNP | exonic | TET2 | nonsynonymous SNV | TET2:NM_001127208:exon4:c.G3484A:p.E1162K |
| chr4 | 106162571 | 106162571 | A | C | m_13_D | TET2 | Missense_Mutation | NM_001127208 | exon4 | c.A3485C | p.E1162A | SNP | exonic | TET2 | nonsynonymous SNV | TET2:NM_001127208:exon4:c.A3485C:p.E1162A |
| chr4 | 106162573 | 106162573 | A | C | m_13_D | TET2 | Missense_Mutation | NM_001127208 | exon4 | c.A3487C | p.I1163L | SNP | exonic | TET2 | nonsynonymous SNV | TET2:NM_001127208:exon4:c.A3487C:p.I1163L |
| chr4 | 106180893 | 106180893 | G | T | m_17_D | TET2 | Missense_Mutation | NM_001127208 | exon7 | c.G3921T | p.R1307S | SNP | exonic | TET2 | nonsynonymous SNV | TET2:NM_001127208:exon7:c.G3921T:p.R1307S |
| chr4 | 106196213 | 106196213 | C | T | m_17_D | TET2 | Nonsense_Mutation | NM_001127208 | exon11 | c.C4546T | p.R1516X | SNP | exonic | TET2 | stopgain | TET2:NM_001127208:exon11:c.C4546T:p.R1516X |
| chr4 | 106164068 | 106164068 | G | A | m_24_D | TET2 | Missense_Mutation | NM_001127208 | exon5 | c.G3578A | p.C1193Y | SNP | exonic | TET2 | nonsynonymous SNV | TET2:NM_001127208:exon5:c.G3578A:p.C1193Y |
| chr4 | 106193748 | 106193748 | C | T | m_24_D | TET2 | Nonsense_Mutation | NM_001127208 | exon10 | c.C4210T | p.R1404X | SNP | exonic | TET2 | stopgain | TET2:NM_001127208:exon10:c.C4210T:p.R1404X |
| chr4 | 106164061 | 106164061 | C | T | m_4_D | TET2 | Nonsense_Mutation | NM_001127208 | exon5 | c.C3571T | p.Q1191X | SNP | exonic | TET2 | stopgain | TET2:NM_001127208:exon5:c.C3571T:p.Q1191X |

| cytoBand | SIFT_score | SIFT_pred | Polyphen2_HDIV_score | Polyphen2_HDIV_pred | Polyphen2_HVAR_score | Polyphen2_HVAR_pred | LRT_score | LRT_pred | MutationTaster_score | MutationTaster_pred | MutationAssessor_score | MutationAssessor_pred | FATHMM_score | FATHMM_pred | PROVEAN_score | PROVEAN_pred | VEST3_score | CADD_raw |
|---|---|---|---|---|---|---|---|---|---|---|---|---|---|---|---|---|---|---|
| 4q24 | 0.112 | T | 0.944 | P | 0.776 | P | . | . | 1 | D | 1.265 | L | 1.45 | T | -4.97 | D | 0.77 | 3.813 |
| 4q24 | 0.675 | T | 0.052 | B | 0.027 | B | . | . | 1 | D | -1.365 | N | 1.58 | T | 1.35 | N | 0.418 | 0.775 |
| 4q24 | 0.148 | T | 0.066 | B | 0.067 | B | . | . | 1 | D | 0.805 | L | 1.22 | T | -2.14 | N | 0.214 | 2.221 |
| 4q24 | 0.022 | D | 0.291 | B | 0.157 | B | . | . | 1 | D | 1.04 | L | 1.12 | T | -3.1 | D | 0.368 | 3.156 |
| 4q24 | 0.007 | D | 0.82 | P | 0.41 | B | . | . | 1 | D | 1.73 | L | 1.11 | T | -4.73 | D | 0.601 | 3.220 |
| 4q24 | 0.314 | T | 0.013 | B | 0.028 | B | . | . | 1 | D | -0.05 | N | 1.35 | T | -0.3 | N | 0.217 | 1.471 |
| 4q24 | 0.001 | D | 1.0 | D | 0.997 | D | . | . | 1 | D | 2.74 | M | 2.48 | T | -5.36 | D | 0.954 | 5.530 |
| 4q24 | . | . | . | . | . | . | . | . | 1 | D | . | . | . | . | . | . | . | 12.788 |
| 4q24 | 0.0 | D | 1.0 | D | 0.999 | D | . | . | 1 | D | 2.38 | M | 0.8 | T | -10.54 | D | 0.958 | 5.981 |
| 4q24 | . | . | . | . | . | . | . | . | 1 | A | . | . | . | . | . | . | . | 15.032 |
| 4q24 | . | . | . | . | . | . | . | . | 1 | A | . | . | . | . | . | . | . | 13.110 |

| CADD_phred | DANN_score | fathmm-MKL coding score | fathmm-MKL coding pred | MetaSVM score | MetaSVM pred | MetaLR score | MetaLR pred | integrated fitCons score | integrated confidence value | GERP++ RS | phyloP7way vertebrate | phyloP20way mammalian | phastCons7way vertebrate | phastCons20way mammalian | SiPhy 29way logOdds |
|---|---|---|---|---|---|---|---|---|---|---|---|---|---|---|---|
| 23.4 | 0.995 | 0.987 | D | -1.036 | T | 0.120 | T | 0.651 | 0 | 5.23 | -0.023 | 0.079 | 0.999 | 1.000 | 17.699 |
| 9.309 | 0.868 | 0.803 | D | -1.017 | T | 0.020 | T | 0.651 | 0 | 4.87 | 0.048 | 0.086 | 1.000 | 1.000 | 10.408 |
| 17.65 | 0.993 | 0.983 | D | -1.098 | T | 0.066 | T | 0.651 | 0 | 4.86 | 0.069 | 0.237 | 1.000 | 1.000 | 12.520 |
| 22.6 | 0.998 | 0.919 | D | -1.050 | T | 0.082 | T | 0.651 | 0 | 4.42 | 0.034 | 0.154 | 0.998 | 0.999 | 9.024 |
| 22.7 | 0.994 | 0.964 | D | -0.959 | T | 0.125 | T | 0.651 | 0 | 4.98 | 0.042 | 0.157 | 0.997 | 0.998 | 13.443 |
| 13.16 | 0.953 | 0.877 | D | -1.051 | T | 0.038 | T | 0.651 | 0 | 3.64 | 0.075 | 0.157 | 0.998 | 0.999 | 9.812 |
| 26.3 | 0.997 | 0.829 | D | -1.104 | T | 0.090 | T | 0.707 | 0 | 2.46 | -0.011 | 0.135 | 0.999 | 0.998 | 4.517 |
| 40 | 0.997 | 0.740 | D | . | . | . | . | 0.672 | 0 | 3.36 | 0.871 | 0.892 | 0.440 | 0.913 | 8.250 |
| 27.8 | 0.997 | 0.992 | D | -1.229 | T | 0.046 | T | 0.707 | 0 | 5.81 | 0.917 | 1.048 | 1.000 | 1.000 | 20.078 |
| 49 | 0.998 | 0.977 | D | . | . | . | . | 0.563 | 0 | 5.11 | 0.871 | 0.935 | 0.987 | 0.995 | 19.089 |
| 41 | 0.998 | 0.986 | D | . | . | . | . | 0.707 | 0 | 5.91 | 0.871 | 0.935 | 1.000 | 1.000 | 20.298 |

**Supplementary Table 1. Top 100 mutated genes in HMCL.**
The table depicts the Var I Decrypt output with the top 100 mutated genes in HMCLs. Samples are highlighted in yellow, variant types and frequency are indicated in grey.

| Gene | Mutation rate (%) | Total number of variants | Frameshift deletion | Frameshift insertion | Non-frameshift deletion | Non-frameshift insertion | Non-synonymous SNV | STOP gain | STOP loss | unknown | AMO1 | JJN3 | L363 | LP1 | Lopra | MM1S | MOLP2 | MOLP8 | OPM2 | RPMI | SKMM2 | XG11 | XG12 | XG13 | XG16 | XG19 | XG1 | XG20 | XG23 | XG24 | XG25 | XG26 | XG27 | XG28 | XG29 | XG2 | XG30 | XG5 | XG6 | XG7 |
|---|---|---|---|---|---|---|---|---|---|---|---|---|---|---|---|---|---|---|---|---|---|---|---|---|---|---|---|---|---|---|---|---|---|---|---|---|---|---|---|---|
| MAP3K1 | 100 | 6 | 0 | 0 | 3 | 1 | 2 | 0 | 0 | 0 | 1 | 1 | 1 | 2 | 1 | 1 | 1 | 2 | 1 | 1 | 1 | 1 | 1 | 1 | 1 | 1 | 1 | 1 | 1 | 1 | 2 | 1 | 1 | 1 | 1 | 1 | 2 | 2 | 2 | 2 |
| CEP89 | 100 | 4 | 0 | 0 | 0 | 0 | 4 | 0 | 0 | 0 | 2 | 3 | 3 | 3 | 3 | 2 | 2 | 3 | 2 | 3 | 3 | 3 | 2 | 2 | 2 | 2 | 3 | 3 | 2 | 2 | 3 | 2 | 3 | 3 | 3 | 3 | 2 | 3 | 1 | 3 |
| RECQL4 | 100 | 2 | 1 | 0 | 0 | 0 | 1 | 0 | 0 | 0 | 1 | 1 | 1 | 1 | 1 | 1 | 1 | 1 | 1 | 1 | 1 | 1 | 1 | 1 | 2 | 1 | 1 | 1 | 1 | 1 | 1 | 1 | 1 | 1 | 1 | 1 | 1 | 1 | 1 | 1 |
| CREB3L1 | 100 | 2 | 0 | 2 | 0 | 0 | 0 | 0 | 0 | 0 | 1 | 1 | 1 | 1 | 2 | 1 | 1 | 1 | 1 | 1 | 1 | 1 | 1 | 1 | 1 | 1 | 1 | 1 | 1 | 1 | 1 | 1 | 1 | 2 | 2 | 1 | 1 | 1 | 2 | 1 |
| KMT2D | 96.7 | 11 | 0 | 0 | 0 | 0 | 9 | 2 | 0 | 0 | 1 | 1 | 1 | 2 | 1 | 1 | 0 | 2 | 3 | 2 | 1 | 2 | 1 | 1 | 1 | 1 | 1 | 1 | 1 | 1 | 1 | 1 | 2 | 5 | 2 | 2 | 1 | 1 | 1 | 1 |
| ZNF384 | 96.7 | 4 | 0 | 0 | 3 | 1 | 0 | 0 | 0 | 0 | 1 | 1 | 1 | 1 | 1 | 1 | 1 | 2 | 1 | 1 | 1 | 1 | 0 | 1 | 1 | 1 | 1 | 1 | 2 | 1 | 1 | 2 | 2 | 1 | 1 | 1 | 1 | 1 | 1 | 1 |
| MAML2 | 93.3 | 3 | 0 | 0 | 1 | 0 | 2 | 0 | 0 | 0 | 1 | 1 | 1 | 1 | 1 | 1 | 1 | 2 | 1 | 1 | 1 | 1 | 1 | 1 | 1 | 1 | 1 | 0 | 1 | 1 | 1 | 1 | 1 | 1 | 1 | 0 | 1 | 1 | 2 | 1 |
| CREB3L2 | 90 | 1 | 0 | 0 | 1 | 0 | 0 | 0 | 0 | 0 | 1 | 1 | 1 | 1 | 1 | 1 | 1 | 1 | 1 | 1 | 1 | 0 | 1 | 1 | 1 | 1 | 1 | 1 | 1 | 1 | 1 | 0 | 1 | 1 | 1 | 1 | 0 | 1 | 1 | 1 |
| SUFU | 83.3 | 7 | 1 | 2 | 0 | 0 | 4 | 0 | 0 | 0 | 2 | 4 | 2 | 0 | 2 | 0 | 0 | 0 | 3 | 5 | 2 | 1 | 4 | 3 | 2 | 3 | 2 | 3 | 2 | 3 | 2 | 3 | 2 | 3 | 0 | 1 | 1 | 1 | 2 | 4 |
| BRD3 | 76.7 | 7 | 0 | 0 | 0 | 0 | 7 | 0 | 0 | 0 | 2 | 1 | 1 | 3 | 0 | 1 | 0 | 1 | 3 | 0 | 2 | 1 | 2 | 1 | 1 | 2 | 1 | 0 | 2 | 1 | 1 | 2 | 2 | 0 | 1 | 2 | 0 | 0 | 2 | 1 |
| PRDM2 | 70 | 5 | 0 | 0 | 1 | 3 | 1 | 0 | 0 | 0 | 1 | 1 | 2 | 1 | 0 | 1 | 1 | 1 | 2 | 1 | 1 | 0 | 1 | 2 | 0 | 1 | 2 | 1 | 0 | 0 | 1 | 2 | 0 | 0 | 0 | 0 | 1 | 1 | 1 | 1 |
| AKAP9 | 63.3 | 3 | 0 | 0 | 0 | 1 | 2 | 0 | 0 | 0 | 1 | 0 | 0 | 0 | 0 | 1 | 1 | 0 | 0 | 0 | 1 | 1 | 1 | 1 | 1 | 1 | 1 | 0 | 1 | 1 | 1 | 0 | 0 | 1 | 2 | 1 | 0 | 1 | 1 | 1 |
| FUS | 63.3 | 2 | 0 | 0 | 1 | 0 | 1 | 0 | 0 | 0 | 1 | 1 | 1 | 0 | 1 | 1 | 0 | 0 | 1 | 1 | 1 | 1 | 1 | 0 | 0 | 0 | 1 | 0 | 1 | 1 | 1 | 1 | 0 | 0 | 0 | 1 | 1 | 0 | 1 | 1 |
| SETD1B | 56.7 | 10 | 0 | 0 | 0 | 0 | 10 | 0 | 0 | 0 | 2 | 1 | 0 | 1 | 1 | 0 | 0 | 0 | 2 | 0 | 1 | 1 | 0 | 1 | 1 | 0 | 2 | 2 | 0 | 0 | 0 | 0 | 2 | 2 | 1 | 1 | 0 | 1 | 4 | 0 |
| FGFR4 | 56.7 | 3 | 0 | 0 | 0 | 0 | 3 | 0 | 0 | 0 | 2 | 1 | 0 | 0 | 0 | 0 | 1 | 1 | 0 | 0 | 1 | 2 | 2 | 2 | 2 | 2 | 0 | 1 | 1 | 3 | 1 | 1 | 0 | 0 | 0 | 1 | 0 | 0 | 0 | 1 |
| HOXC13 | 50 | 1 | 0 | 0 | 0 | 0 | 1 | 0 | 0 | 0 | 1 | 1 | 0 | 1 | 1 | 1 | 0 | 1 | 1 | 0 | 0 | 1 | 0 | 0 | 1 | 0 | 0 | 0 | 1 | 1 | 0 | 0 | 0 | 0 | 1 | 0 | 1 | 0 | 1 | 1 |
| TP53 | 46.7 | 14 | 0 | 0 | 0 | 0 | 13 | 1 | 0 | 0 | 0 | 0 | 1 | 1 | 0 | 0 | 1 | 0 | 1 | 1 | 1 | 1 | 0 | 1 | 1 | 1 | 1 | 0 | 0 | 0 | 0 | 0 | 0 | 0 | 0 | 2 | 1 | 1 | 0 | 0 |
| FANCD2 | 46.7 | 4 | 0 | 0 | 0 | 0 | 4 | 0 | 0 | 0 | 1 | 1 | 1 | 1 | 0 | 0 | 0 | 0 | 0 | 0 | 0 | 1 | 1 | 0 | 0 | 0 | 0 | 1 | 0 | 0 | 2 | 1 | 1 | 1 | 1 | 0 | 0 | 0 | 2 | 1 |
| SMC1A | 46.7 | 4 | 0 | 0 | 0 | 0 | 4 | 0 | 0 | 0 | 0 | 1 | 0 | 3 | 0 | 1 | 0 | 0 | 2 | 1 | 2 | 1 | 2 | 2 | 0 | 0 | 0 | 0 | 1 | 0 | 1 | 0 | 0 | 0 | 0 | 1 | 0 | 0 | 2 | 1 |
| MN1 | 43.3 | 11 | 0 | 0 | 0 | 9 | 0 | 2 | 0 | 0 | 0 | 0 | 0 | 0 | 2 | 2 | 0 | 0 | 0 | 2 | 0 | 0 | 0 | 0 | 1 | 0 | 1 | 1 | 0 | 0 | 2 | 0 | 1 | 3 | 2 | 1 | 0 | 0 | 2 | 2 |
| MDS2 | 43.3 | 5 | 0 | 0 | 0 | 4 | 0 | 1 | 0 | 0 | 1 | 0 | 0 | 0 | 1 | 0 | 0 | 2 | 0 | 0 | 1 | 1 | 1 | 0 | 0 | 0 | 1 | 0 | 0 | 0 | 1 | 0 | 2 | 2 | 0 | 1 | 1 | 0 | 0 | 2 |
| BRD4 | 43.3 | 5 | 0 | 0 | 0 | 0 | 5 | 0 | 0 | 0 | 0 | 0 | 1 | 0 | 1 | 3 | 0 | 0 | 2 | 0 | 0 | 2 | 0 | 1 | 0 | 2 | 0 | 1 | 1 | 1 | 0 | 1 | 0 | 0 | 0 | 0 | 0 | 0 | 1 | 1 |
| CDK12 | 43.3 | 4 | 0 | 0 | 0 | 0 | 3 | 1 | 0 | 0 | 1 | 1 | 2 | 1 | 0 | 0 | 0 | 0 | 0 | 0 | 3 | 2 | 1 | 0 | 2 | 0 | 0 | 0 | 0 | 1 | 3 | 1 | 0 | 0 | 0 | 1 | 0 | 1 | 0 | 0 |
| CNOT3 | 43.3 | 4 | 0 | 0 | 0 | 0 | 4 | 0 | 0 | 0 | 0 | 0 | 2 | 3 | 0 | 0 | 0 | 1 | 2 | 0 | 3 | 0 | 1 | 1 | 1 | 0 | 1 | 0 | 0 | 2 | 1 | 1 | 0 | 0 | 0 | 0 | 0 | 0 | 1 | 0 |
| KIAA1549 | 43.3 | 3 | 0 | 0 | 0 | 0 | 3 | 0 | 0 | 0 | 0 | 0 | 0 | 0 | 0 | 0 | 1 | 1 | 2 | 1 | 1 | 1 | 0 | 0 | 1 | 1 | 1 | 0 | 0 | 0 | 0 | 1 | 0 | 0 | 0 | 1 | 1 | 1 | 0 | 0 |
| FNBP1 | 43.3 | 1 | 0 | 0 | 0 | 0 | 1 | 0 | 0 | 0 | 0 | 0 | 1 | 1 | 0 | 0 | 0 | 0 | 0 | 1 | 1 | 1 | 1 | 0 | 1 | 1 | 1 | 1 | 0 | 0 | 0 | 0 | 0 | 0 | 0 | 1 | 1 | 0 | 0 | 1 |
| KRAS | 40 | 11 | 0 | 0 | 1 | 0 | 10 | 0 | 0 | 0 | 1 | 0 | 0 | 0 | 0 | 1 | 0 | 1 | 0 | 1 | 0 | 1 | 0 | 0 | 0 | 0 | 0 | 1 | 0 | 0 | 2 | 0 | 1 | 0 | 2 | 1 | 0 | 0 | 2 | 2 |
| LRP1B | 40 | 8 | 0 | 0 | 0 | 0 | 8 | 0 | 0 | 0 | 2 | 0 | 1 | 1 | 0 | 0 | 0 | 0 | 1 | 0 | 1 | 0 | 1 | 0 | 1 | 0 | 2 | 0 | 1 | 1 | 0 | 2 | 0 | 0 | 0 | 0 | 0 | 0 | 1 | 0 |
| RET | 40 | 2 | 1 | 0 | 0 | 0 | 1 | 0 | 0 | 0 | 0 | 0 | 0 | 0 | 1 | 0 | 0 | 0 | 0 | 0 | 0 | 1 | 0 | 1 | 1 | 0 | 0 | 1 | 1 | 1 | 1 | 1 | 0 | 0 | 1 | 0 | 1 | 0 | 0 | 1 |
| MUC16 | 36.7 | 16 | 0 | 1 | 0 | 0 | 15 | 0 | 0 | 0 | 0 | 0 | 1 | 0 | 2 | 0 | 0 | 0 | 1 | 0 | 0 | 2 | 2 | 0 | 0 | 1 | 1 | 0 | 1 | 2 | 0 | 0 | 0 | 0 | 0 | 0 | 2 | 0 | 2 | 0 |
| ARID1B | 36.7 | 10 | 0 | 0 | 2 | 6 | 2 | 0 | 0 | 0 | 0 | 0 | 0 | 1 | 0 | 0 | 0 | 0 | 0 | 3 | 0 | 1 | 0 | 0 | 0 | 0 | 1 | 2 | 0 | 0 | 1 | 2 | 0 | 1 | 0 | 0 | 1 | 0 | 1 | 1 |
| ETV5 | 36.7 | 5 | 0 | 1 | 0 | 0 | 4 | 0 | 0 | 0 | 1 | 0 | 1 | 0 | 0 | 0 | 0 | 0 | 1 | 0 | 0 | 0 | 1 | 1 | 2 | 0 | 0 | 2 | 1 | 1 | 0 | 0 | 0 | 0 | 0 | 1 | 0 | 0 | 1 | 0 |
| PABPC1 | 36.7 | 5 | 0 | 0 | 0 | 0 | 5 | 0 | 0 | 0 | 0 | 0 | 1 | 2 | 0 | 4 | 1 | 3 | 2 | 3 | 0 | 3 | 0 | 2 | 2 | 0 | 0 | 0 | 0 | 0 | 0 | 0 | 0 | 0 | 0 | 0 | 2 | 0 | 0 | 0 |
| SIRPA | 36.7 | 3 | 0 | 0 | 1 | 0 | 2 | 0 | 0 | 0 | 3 | 0 | 0 | 2 | 3 | 0 | 2 | 0 | 2 | 0 | 0 | 0 | 0 | 0 | 0 | 2 | 2 | 0 | 0 | 0 | 2 | 2 | 0 | 0 | 0 | 0 | 0 | 2 | 0 | 3 |
| ATM | 33.3 | 11 | 0 | 0 | 0 | 0 | 10 | 1 | 0 | 0 | 1 | 0 | 0 | 0 | 1 | 0 | 0 | 0 | 0 | 0 | 0 | 1 | 0 | 0 | 0 | 2 | 0 | 1 | 1 | 1 | 0 | 0 | 0 | 0 | 1 | 0 | 2 | 0 | 1 | 0 |
| AFDN | 33.3 | 7 | 0 | 0 | 0 | 0 | 7 | 0 | 0 | 0 | 0 | 0 | 1 | 1 | 0 | 0 | 0 | 0 | 0 | 1 | 0 | 1 | 0 | 0 | 0 | 0 | 1 | 1 | 0 | 0 | 2 | 0 | 1 | 0 | 0 | 0 | 0 | 0 | 1 | 2 |
| CIC | 33.3 | 7 | 0 | 1 | 0 | 1 | 5 | 0 | 0 | 0 | 0 | 0 | 0 | 0 | 0 | 1 | 0 | 2 | 0 | 1 | 0 | 0 | 0 | 0 | 0 | 1 | 0 | 0 | 1 | 1 | 0 | 1 | 0 | 1 | 0 | 0 | 1 | 0 | 0 | 1 |
| PER1 | 33.3 | 6 | 0 | 0 | 0 | 0 | 6 | 0 | 0 | 0 | 1 | 1 | 0 | 0 | 1 | 0 | 0 | 1 | 0 | 0 | 0 | 1 | 1 | 0 | 0 | 0 | 0 | 1 | 2 | 2 | 0 | 0 | 0 | 0 | 1 | 0 | 0 | 0 | 0 | 0 |
| COL1A1 | 33.3 | 5 | 0 | 1 | 0 | 0 | 4 | 0 | 0 | 0 | 0 | 1 | 2 | 1 | 1 | 0 | 0 | 0 | 2 | 0 | 0 | 1 | 0 | 1 | 0 | 0 | 0 | 0 | 0 | 1 | 0 | 0 | 0 | 0 | 1 | 0 | 0 | 0 | 0 | 1 |
| POLG | 33.3 | 5 | 0 | 0 | 1 | 1 | 3 | 0 | 0 | 0 | 0 | 0 | 0 | 0 | 0 | 0 | 1 | 0 | 0 | 0 | 0 | 0 | 0 | 1 | 1 | 0 | 0 | 1 | 2 | 2 | 2 | 1 | 0 | 1 | 0 | 0 | 0 | 0 | 0 | 2 |
| NRAS | 30 | 6 | 0 | 0 | 0 | 0 | 6 | 0 | 0 | 0 | 0 | 0 | 1 | 0 | 1 | 0 | 0 | 1 | 0 | 0 | 0 | 0 | 1 | 0 | 1 | 0 | 1 | 0 | 1 | 1 | 0 | 1 | 0 | 0 | 0 | 0 | 0 | 0 | 0 | 0 |
| KAT6B | 30 | 1 | 0 | 0 | 1 | 0 | 0 | 0 | 0 | 0 | 1 | 0 | 0 | 0 | 0 | 1 | 1 | 0 | 0 | 1 | 1 | 0 | 1 | 0 | 0 | 0 | 0 | 0 | 1 | 1 | 0 | 0 | 0 | 0 | 1 | 0 | 0 | 0 | 0 | 0 |
| ZFHX3 | 26.7 | 10 | 0 | 0 | 3 | 1 | 6 | 0 | 0 | 0 | 1 | 4 | 0 | 2 | 0 | 0 | 0 | 0 | 1 | 2 | 2 | 0 | 1 | 0 | 0 | 0 | 0 | 1 | 0 | 0 | 0 | 0 | 0 | 0 | 0 | 0 | 0 | 0 | 0 | 0 |
| PTPRD | 26.7 | 7 | 0 | 1 | 0 | 0 | 5 | 1 | 0 | 0 | 1 | 0 | 0 | 0 | 0 | 2 | 4 | 0 | 0 | 0 | 0 | 0 | 1 | 1 | 0 | 0 | 1 | 0 | 1 | 2 | 0 | 0 | 0 | 0 | 0 | 0 | 0 | 0 | 0 | 0 |
| MRTFA | 26.7 | 7 | 0 | 1 | 1 | 1 | 4 | 0 | 0 | 0 | 0 | 0 | 0 | 0 | 0 | 1 | 0 | 1 | 0 | 0 | 0 | 1 | 2 | 1 | 0 | 0 | 0 | 0 | 1 | 1 | 1 | 0 | 0 | 0 | 0 | 0 | 0 | 0 | 0 | 0 |
| AFF3 | 26.7 | 4 | 0 | 0 | 2 | 0 | 2 | 0 | 0 | 0 | 1 | 0 | 2 | 0 | 0 | 0 | 3 | 0 | 0 | 0 | 0 | 0 | 0 | 0 | 0 | 2 | 0 | 0 | 0 | 1 | 2 | 0 | 0 | 0 | 0 | 0 | 2 | 0 | 0 | 2 |
| IGF2BP2 | 26.7 | 1 | 0 | 0 | 0 | 0 | 1 | 0 | 0 | 0 | 0 | 0 | 0 | 0 | 0 | 0 | 0 | 0 | 1 | 0 | 1 | 1 | 1 | 0 | 0 | 1 | 0 | 0 | 0 | 1 | 0 | 0 | 0 | 0 | 0 | 0 | 0 | 0 | 1 | 1 |
| TENT5C | 23.3 | 26 | 1 | 5 | 0 | 0 | 20 | 0 | 0 | 0 | 0 | 0 | 0 | 0 | 0 | 1 | 0 | 2 | 1 | 0 | 0 | 0 | 0 | 0 | 0 | 0 | 0 | 8 | 0 | 0 | 12 | 0 | 0 | 0 | 1 | 0 | 0 | 0 | 0 | 6 |
| TNC | 23.3 | 7 | 0 | 0 | 0 | 0 | 7 | 0 | 0 | 0 | 1 | 0 | 0 | 0 | 0 | 0 | 1 | 1 | 0 | 0 | 0 | 0 | 1 | 0 | 0 | 1 | 1 | 0 | 0 | 0 | 0 | 0 | 0 | 0 | 0 | 0 | 0 | 0 | 0 | 1 |
| FAT4 | 23.3 | 6 | 0 | 0 | 0 | 0 | 6 | 0 | 0 | 0 | 1 | 0 | 0 | 0 | 0 | 0 | 1 | 0 | 1 | 0 | 0 | 0 | 0 | 0 | 0 | 0 | 0 | 0 | 0 | 0 | 1 | 0 | 1 | 0 | 0 | 0 | 0 | 0 | 1 | 1 |
| GNAS | 23.3 | 6 | 0 | 1 | 0 | 0 | 5 | 0 | 0 | 0 | 0 | 0 | 0 | 0 | 0 | 0 | 0 | 1 | 1 | 0 | 0 | 1 | 0 | 0 | 1 | 0 | 0 | 0 | 0 | 0 | 1 | 1 | 0 | 1 | 0 | 0 | 0 | 0 | 0 | 0 |
| CREBBP | 23.3 | 4 | 0 | 0 | 0 | 0 | 4 | 0 | 0 | 0 | 1 | 0 | 0 | 0 | 0 | 0 | 1 | 0 | 0 | 0 | 0 | 0 | 0 | 0 | 0 | 1 | 0 | 1 | 0 | 0 | 1 | 1 | 0 | 0 | 0 | 0 | 0 | 0 | 2 | 0 |
| RUNX1 | 20 | 7 | 0 | 0 | 0 | 0 | 7 | 0 | 0 | 0 | 0 | 0 | 0 | 1 | 4 | 0 | 0 | 0 | 0 | 0 | 0 | 0 | 0 | 0 | 0 | 0 | 1 | 0 | 2 | 0 | 0 | 0 | 1 | 0 | 0 | 0 | 3 | 0 | 0 | 0 |
| CSMD3 | 20 | 6 | 0 | 0 | 0 | 0 | 6 | 0 | 0 | 0 | 0 | 1 | 0 | 0 | 1 | 0 | 0 | 0 | 0 | 0 | 0 | 0 | 0 | 0 | 0 | 0 | 2 | 0 | 1 | 1 | 0 | 0 | 1 | 0 | 0 | 0 | 0 | 0 | 0 | 0 |
| NUP214 | 20 | 6 | 1 | 0 | 0 | 0 | 5 | 0 | 0 | 0 | 0 | 0 | 0 | 1 | 0 | 0 | 0 | 0 | 0 | 1 | 0 | 1 | 0 | 1 | 0 | 0 | 0 | 0 | 0 | 0 | 0 | 0 | 1 | 0 | 0 | 0 | 1 | 0 | 0 | 0 |
| SPEN | 20 | 5 | 0 | 0 | 0 | 0 | 5 | 0 | 0 | 0 | 1 | 0 | 0 | 0 | 0 | 0 | 0 | 0 | 0 | 1 | 0 | 0 | 0 | 0 | 1 | 0 | 0 | 0 | 0 | 0 | 1 | 0 | 1 | 0 | 0 | 0 | 0 | 0 | 0 | 1 |
| HEY1 | 20 | 4 | 2 | 0 | 0 | 1 | 1 | 0 | 0 | 0 | 0 | 0 | 0 | 1 | 0 | 0 | 0 | 0 | 0 | 2 | 0 | 0 | 0 | 0 | 1 | 0 | 0 | 0 | 3 | 1 | 0 | 0 | 0 | 1 | 0 | 0 | 0 | 0 | 0 | 0 |
| NACA | 20 | 3 | 0 | 1 | 0 | 0 | 2 | 0 | 0 | 0 | 0 | 1 | 0 | 0 | 1 | 0 | 0 | 0 | 0 | 0 | 0 | 1 | 1 | 0 | 0 | 0 | 0 | 1 | 0 | 0 | 0 | 0 | 0 | 0 | 0 | 0 | 0 | 0 | 0 | 1 |
| ELL | 20 | 2 | 0 | 0 | 0 | 0 | 2 | 0 | 0 | 0 | 0 | 1 | 0 | 0 | 1 | 0 | 0 | 0 | 1 | 0 | 0 | 0 | 1 | 0 | 0 | 1 | 0 | 0 | 0 | 1 | 0 | 0 | 0 | 0 | 0 | 0 | 0 | 0 | 0 | 0 |
| FLT4 | 20 | 2 | 0 | 0 | 0 | 0 | 2 | 0 | 0 | 0 | 0 | 0 | 0 | 2 | 0 | 0 | 0 | 0 | 0 | 0 | 1 | 0 | 0 | 0 | 0 | 0 | 1 | 0 | 0 | 0 | 1 | 0 | 1 | 0 | 0 | 0 | 0 | 0 | 0 | 1 |
| ARID1A | 16.7 | 5 | 0 | 0 | 0 | 0 | 5 | 0 | 0 | 0 | 1 | 0 | 1 | 0 | 0 | 0 | 0 | 0 | 1 | 0 | 0 | 0 | 0 | 0 | 0 | 1 | 0 | 0 | 0 | 0 | 1 | 0 | 0 | 0 | 0 | 0 | 0 | 0 | 0 | 0 |
| BCL9L | 16.7 | 5 | 0 | 1 | 0 | 0 | 4 | 0 | 0 | 0 | 0 | 0 | 0 | 0 | 0 | 0 | 0 | 1 | 0 | 0 | 1 | 1 | 0 | 0 | 0 | 0 | 0 | 0 | 0 | 1 | 0 | 1 | 0 | 0 | 0 | 0 | 0 | 0 | 0 | 0 |
| BIRC6 | 16.7 | 4 | 0 | 0 | 0 | 0 | 4 | 0 | 0 | 0 | 0 | 1 | 0 | 0 | 0 | 0 | 0 | 0 | 0 | 0 | 0 | 0 | 0 | 0 | 0 | 0 | 0 | 0 | 0 | 0 | 1 | 0 | 0 | 0 | 0 | 1 | 0 | 1 | 0 | 1 |
| TRIP11 | 16.7 | 3 | 0 | 0 | 0 | 0 | 3 | 0 | 0 | 0 | 0 | 0 | 0 | 0 | 1 | 0 | 1 | 0 | 0 | 0 | 0 | 0 | 0 | 0 | 0 | 1 | 0 | 0 | 0 | 0 | 1 | 0 | 0 | 0 | 0 | 0 | 0 | 0 | 0 | 1 |
| PMS1 | 16.7 | 3 | 0 | 1 | 0 | 0 | 2 | 0 | 0 | 0 | 0 | 0 | 0 | 0 | 0 | 0 | 0 | 0 | 1 | 0 | 1 | 0 | 0 | 0 | 0 | 1 | 0 | 0 | 0 | 0 | 1 | 0 | 0 | 0 | 0 | 0 | 0 | 0 | 0 | 1 |
| ANK1 | 16.7 | 3 | 0 | 0 | 0 | 0 | 3 | 0 | 0 | 0 | 0 | 0 | 0 | 0 | 0 | 0 | 0 | 0 | 0 | 0 | 0 | 0 | 0 | 1 | 0 | 0 | 0 | 0 | 0 | 0 | 0 | 0 | 1 | 1 | 1 | 1 | 0 | 0 | 0 | 0 |
| MNX1 | 16.7 | 2 | 0 | 0 | 1 | 1 | 0 | 0 | 0 | 0 | 0 | 0 | 0 | 0 | 0 | 1 | 0 | 0 | 0 | 0 | 0 | 0 | 0 | 1 | 0 | 0 | 1 | 0 | 0 | 0 | 0 | 0 | 0 | 0 | 1 | 1 | 0 | 0 | 0 | 0 |
| CCND1 | 13.3 | 6 | 0 | 0 | 0 | 0 | 6 | 0 | 0 | 0 | 0 | 0 | 0 | 1 | 0 | 0 | 0 | 0 | 0 | 0 | 0 | 0 | 0 | 0 | 0 | 0 | 0 | 0 | 1 | 1 | 0 | 0 | 0 | 0 | 0 | 0 | 0 | 4 | 0 | 0 |
| NCOA2 | 13.3 | 5 | 0 | 0 | 0 | 0 | 4 | 1 | 0 | 0 | 0 | 3 | 0 | 0 | 0 | 0 | 0 | 0 | 0 | 0 | 0 | 0 | 0 | 0 | 0 | 0 | 0 | 1 | 1 | 1 | 0 | 0 | 0 | 0 | 0 | 0 | 0 | 0 | 0 | 0 |
| IRS4 | 13.3 | 5 | 0 | 0 | 0 | 0 | 5 | 0 | 0 | 0 | 0 | 0 | 2 | 1 | 0 | 0 | 0 | 0 | 0 | 0 | 0 | 0 | 0 | 0 | 0 | 0 | 0 | 0 | 0 | 0 | 0 | 0 | 0 | 1 | 0 | 0 | 1 | 0 | 0 | 0 |
| EGFR | 13.3 | 4 | 0 | 0 | 0 | 0 | 4 | 0 | 0 | 0 | 1 | 0 | 0 | 0 | 1 | 1 | 0 | 0 | 0 | 1 | 0 | 0 | 0 | 0 | 0 | 0 | 0 | 0 | 0 | 0 | 0 | 0 | 0 | 0 | 0 | 0 | 0 | 0 | 0 | 0 |
| FAT3 | 13.3 | 4 | 0 | 0 | 0 | 0 | 4 | 0 | 0 | 0 | 1 | 0 | 0 | 0 | 0 | 0 | 0 | 0 | 1 | 0 | 0 | 0 | 0 | 1 | 0 | 1 | 0 | 0 | 0 | 0 | 0 | 0 | 0 | 0 | 0 | 0 | 0 | 0 | 0 | 0 |
| KIF5B | 13.3 | 4 | 0 | 0 | 0 | 0 | 4 | 0 | 0 | 0 | 0 | 0 | 0 | 0 | 1 | 0 | 0 | 0 | 0 | 0 | 0 | 1 | 0 | 0 | 0 | 0 | 0 | 0 | 0 | 0 | 0 | 0 | 0 | 0 | 0 | 0 | 1 | 1 | 0 | 0 |
| ZNF521 | 13.3 | 4 | 0 | 0 | 0 | 0 | 4 | 0 | 0 | 0 | 0 | 0 | 0 | 0 | 0 | 0 | 1 | 0 | 0 | 1 | 0 | 0 | 0 | 0 | 0 | 1 | 0 | 0 | 0 | 0 | 0 | 0 | 0 | 0 | 0 | 0 | 0 | 0 | 1 | 0 |
| NUP98 | 13.3 | 4 | 0 | 0 | 0 | 0 | 4 | 0 | 0 | 0 | 0 | 0 | 0 | 0 | 0 | 0 | 0 | 0 | 1 | 0 | 2 | 0 | 0 | 0 | 1 | 0 | 0 | 0 | 0 | 1 | 0 | 0 | 0 | 0 | 0 | 0 | 0 | 0 | 0 | 0 |
| TSC2 | 13.3 | 4 | 0 | 0 | 0 | 0 | 4 | 0 | 0 | 0 | 0 | 0 | 0 | 0 | 0 | 0 | 0 | 0 | 0 | 0 | 2 | 0 | 0 | 0 | 0 | 0 | 0 | 0 | 0 | 0 | 1 | 0 | 1 | 0 | 0 | 0 | 0 | 0 | 0 | 1 |
| ASXL1 | 13.3 | 4 | 0 | 0 | 0 | 0 | 4 | 0 | 0 | 0 | 0 | 0 | 0 | 0 | 0 | 0 | 0 | 0 | 0 | 0 | 0 | 1 | 1 | 0 | 0 | 0 | 0 | 0 | 0 | 0 | 0 | 0 | 0 | 0 | 1 | 0 | 1 | 0 | 0 | 0 |
| MYO5A | 13.3 | 4 | 0 | 0 | 0 | 0 | 4 | 0 | 0 | 0 | 0 | 0 | 0 | 0 | 0 | 0 | 0 | 0 | 0 | 0 | 0 | 0 | 1 | 2 | 0 | 0 | 0 | 0 | 1 | 1 | 0 | 0 | 0 | 0 | 0 | 0 | 0 | 0 | 0 | 0 |
| WNK2 | 13.3 | 4 | 0 | 0 | 0 | 0 | 4 | 0 | 0 | 0 | 0 | 0 | 0 | 0 | 0 | 0 | 0 | 0 | 0 | 0 | 0 | 0 | 1 | 0 | 0 | 0 | 1 | 0 | 0 | 0 | 0 | 0 | 0 | 0 | 0 | 1 | 0 | 0 | 2 | 0 |
| PTPRB | 13.3 | 3 | 0 | 0 | 0 | 0 | 3 | 0 | 0 | 0 | 0 | 0 | 1 | 0 | 0 | 0 | 0 | 0 | 0 | 0 | 0 | 0 | 0 | 0 | 0 | 0 | 0 | 0 | 1 | 1 | 0 | 0 | 1 | 0 | 0 | 0 | 0 | 0 | 0 | 0 |
| DCTN1 | 13.3 | 3 | 0 | 0 | 0 | 0 | 3 | 0 | 0 | 0 | 0 | 0 | 0 | 0 | 1 | 0 | 0 | 0 | 0 | 0 | 0 | 0 | 0 | 0 | 0 | 0 | 0 | 0 | 0 | 0 | 1 | 0 | 0 | 0 | 0 | 0 | 0 | 1 | 0 | 1 |
| FAT1 | 13.3 | 3 | 0 | 0 | 0 | 0 | 3 | 0 | 0 | 0 | 0 | 0 | 0 | 0 | 1 | 0 | 0 | 0 | 0 | 0 | 0 | 0 | 0 | 0 | 0 | 0 | 0 | 0 | 1 | 1 | 0 | 0 | 0 | 0 | 0 | 0 | 1 | 0 | 0 | 0 |
| PTEN | 13.3 | 3 | 0 | 0 | 0 | 0 | 3 | 0 | 0 | 0 | 0 | 0 | 0 | 0 | 1 | 1 | 0 | 1 | 0 | 0 | 0 | 0 | 0 | 0 | 0 | 0 | 0 | 0 | 0 | 0 | 0 | 0 | 0 | 0 | 0 | 0 | 1 | 0 | 0 | 0 |
| MAF | 13.3 | 3 | 0 | 0 | 0 | 0 | 3 | 0 | 0 | 0 | 0 | 0 | 0 | 0 | 0 | 1 | 0 | 0 | 0 | 0 | 0 | 0 | 0 | 1 | 0 | 1 | 0 | 0 | 0 | 0 | 0 | 0 | 0 | 0 | 0 | 0 | 0 | 0 | 1 | 0 |
| ARHGEF10 | 13.3 | 3 | 1 | 0 | 0 | 0 | 2 | 0 | 0 | 0 | 0 | 0 | 0 | 0 | 0 | 0 | 0 | 0 | 0 | 0 | 0 | 0 | 0 | 0 | 0 | 0 | 0 | 0 | 0 | 0 | 1 | 0 | 0 | 0 | 0 | 1 | 0 | 0 | 1 | 1 |
| PAX3 | 13.3 | 2 | 0 | 0 | 0 | 0 | 2 | 0 | 0 | 0 | 0 | 1 | 0 | 0 | 0 | 0 | 0 | 0 | 0 | 0 | 0 | 0 | 0 | 0 | 0 | 1 | 0 | 0 | 1 | 1 | 0 | 0 | 0 | 0 | 0 | 0 | 0 | 0 | 0 | 0 |
| PTCH1 | 10 | 7 | 0 | 0 | 0 | 2 | 3 | 2 | 0 | 0 | 0 | 2 | 0 | 0 | 0 | 0 | 0 | 0 | 0 | 3 | 0 | 0 | 0 | 0 | 0 | 0 | 0 | 0 | 0 | 0 | 0 | 0 | 0 | 0 | 0 | 0 | 2 | 0 | 0 | 0 |
| CYLD | 10 | 7 | 1 | 0 | 0 | 0 | 6 | 0 | 0 | 0 | 0 | 0 | 0 | 0 | 0 | 0 | 0 | 0 | 0 | 0 | 0 | 0 | 0 | 0 | 1 | 0 | 0 | 0 | 0 | 0 | 0 | 0 | 0 | 0 | 0 | 0 | 1 | 5 | 0 | 0 |
| NRG1 | 10 | 4 | 0 | 0 | 0 | 0 | 4 | 0 | 0 | 0 | 1 | 0 | 0 | 0 | 1 | 0 | 0 | 0 | 0 | 0 | 0 | 0 | 0 | 0 | 0 | 0 | 0 | 0 | 0 | 0 | 0 | 0 | 0 | 0 | 0 | 2 | 0 | 0 | 0 | 0 |
| STAT3 | 10 | 4 | 1 | 0 | 0 | 0 | 3 | 0 | 0 | 0 | 1 | 0 | 0 | 0 | 0 | 0 | 0 | 0 | 0 | 0 | 0 | 0 | 0 | 1 | 0 | 0 | 0 | 0 | 0 | 0 | 0 | 0 | 0 | 0 | 0 | 0 | 0 | 0 | 2 | 0 |
| RNF213 | 10 | 4 | 0 | 0 | 0 | 0 | 3 | 1 | 0 | 0 | 0 | 0 | 2 | 0 | 0 | 0 | 0 | 0 | 0 | 0 | 0 | 0 | 0 | 0 | 0 | 0 | 0 | 0 | 0 | 0 | 0 | 1 | 0 | 0 | 0 | 1 | 0 | 0 | 0 | 0 |
| TRRAP | 10 | 4 | 0 | 0 | 0 | 0 | 4 | 0 | 0 | 0 | 0 | 0 | 0 | 1 | 0 | 0 | 0 | 0 | 0 | 0 | 0 | 1 | 0 | 0 | 0 | 0 | 2 | 0 | 0 | 0 | 0 | 0 | 0 | 0 | 0 | 0 | 0 | 0 | 0 | 0 |
| GRIN2A | 10 | 4 | 0 | 0 | 0 | 0 | 4 | 0 | 0 | 0 | 0 | 0 | 0 | 0 | 0 | 1 | 0 | 0 | 1 | 0 | 0 | 0 | 0 | 0 | 0 | 2 | 0 | 0 | 0 | 0 | 0 | 0 | 0 | 0 | 0 | 0 | 0 | 0 | 0 | 0 |
| ROBO2 | 10 | 4 | 0 | 0 | 0 | 0 | 4 | 0 | 0 | 0 | 0 | 0 | 0 | 0 | 0 | 1 | 0 | 0 | 0 | 0 | 0 | 0 | 0 | 0 | 0 | 0 | 0 | 0 | 0 | 0 | 0 | 0 | 0 | 0 | 0 | 0 | 1 | 0 | 2 | 0 |
| NCOR1 | 10 | 4 | 0 | 0 | 0 | 0 | 3 | 1 | 0 | 0 | 0 | 0 | 0 | 0 | 0 | 0 | 0 | 0 | 1 | 0 | 1 | 0 | 0 | 0 | 0 | 0 | 0 | 0 | 0 | 0 | 0 | 0 | 0 | 0 | 0 | 0 | 0 | 2 | 0 | 0 |
| PRDM1 | 10 | 4 | 0 | 0 | 0 | 0 | 4 | 0 | 0 | 0 | 0 | 0 | 0 | 0 | 0 | 0 | 0 | 0 | 0 | 0 | 0 | 0 | 0 | 0 | 0 | 0 | 0 | 0 | 1 | 1 | 0 | 0 | 0 | 3 | 0 | 0 | 0 | 0 | 0 | 0 |
| CNBD1 | 10 | 3 | 0 | 0 | 0 | 0 | 1 | 2 | 0 | 0 | 1 | 0 | 0 | 0 | 0 | 0 | 0 | 0 | 0 | 0 | 0 | 1 | 0 | 0 | 0 | 0 | 0 | 0 | 0 | 0 | 0 | 0 | 0 | 1 | 0 | 0 | 0 | 0 | 0 | 0 |
| MSH6 | 10 | 3 | 0 | 0 | 0 | 0 | 3 | 0 | 0 | 0 | 0 | 0 | 1 | 0 | 0 | 0 | 0 | 0 | 1 | 0 | 0 | 0 | 0 | 0 | 0 | 0 | 0 | 1 | 0 | 0 | 0 | 0 | 0 | 0 | 0 | 0 | 0 | 0 | 0 | 0 |
| ERBB3 | 10 | 3 | 0 | 1 | 0 | 0 | 2 | 0 | 0 | 0 | 0 | 0 | 1 | 0 | 1 | 0 | 0 | 0 | 0 | 0 | 0 | 0 | 1 | 0 | 0 | 0 | 0 | 0 | 0 | 0 | 0 | 0 | 0 | 0 | 0 | 0 | 0 | 0 | 0 | 0 |
| USP6 | 10 | 3 | 0 | 0 | 0 | 0 | 3 | 0 | 0 | 0 | 0 | 0 | 2 | 0 | 0 | 0 | 0 | 0 | 0 | 0 | 0 | 0 | 0 | 0 | 0 | 0 | 0 | 0 | 2 | 0 | 0 | 0 | 0 | 0 | 0 | 0 | 1 | 0 | 0 | 0 |
